# Extracellular Vesicles from Stem cells Rescue Cellular Phenotypes and Behavioral Deficits in SHANK3-Associated ASD Neuronal and Mouse Models

**DOI:** 10.1101/2024.08.27.609917

**Authors:** Ashwani Choudhary, Idan Rosh, Yara Hussein, Shai Netser, Aviram Shemen, Tagreed Suliman, Wote Amelo Rike, Lilach Simchi, Boris Shklyar, Ahmad Abu-Akel, Assaf Zinger, Daniel Offen, Shlomo Wagner, Shani Stern

## Abstract

Extracellular vesicles (EVs) are lipid membrane-bound structures that mediate intercellular communication by transferring diverse cargoes, including RNA and proteins. *Shank3*, a synaptic scaffolding protein critical for synapse structure and function, is implicated in autism spectrum disorder (ASD) and Phelan-McDermid Syndrome (PMS). Early hyperexcitability in cortical neurons is a recognized endophenotype in ASD. Here, we investigated EV-mediated effects in the context of *Shank3* deficiency using human iPSC-derived cortical neurons and *Shank3B*-/- mice. Switching EVs between *Shank3* mutant and control neurons revealed that *Shank3* mutant-derived EVs transferred the hyperexcitability and accelerated maturation phenotypes to control neurons. This was driven by enriched synaptic proteins (e.g., *ACTB*, *CFL1*, *AGRN*, *CLSTN1*) in *Shank3* mutant-derived EVs as confirmed by proteomic analysis. Conversely, control EVs failed to rescue mutant phenotypes consistent with their lower enrichment for synaptic proteins and related pathways. Further, EVs from mesenchymal stem cells (MSCs) and healthy donor iPSCs, containing synaptic modulators such as complement proteins (*C1R*, *C1S*), plasticity-related proteins (*MDK*, *IGFBP3*), and homeostatic regulators (*FGF2*, *SFRP1*), rescued the hyperexcitability and normalized the maturation in *Shank3* mutant neurons. Moreover, intranasal administration of iPSC-derived EVs in *Shank3B*-/- mice significantly ameliorated ASD-like behavioral deficits, underscoring their therapeutic potential. Together, these findings reveal a novel EV-mediated mechanism for modulating dysregulated excitability and synaptic maturation, addressing a critical unmet need in ASD and related neurodevelopmental disorders treatment.

## INTRODUCTION

The *Shank3* gene belongs to the family of ProSAP/Shank genes(*1*, *2*) and plays a key role in maintaining synaptic plasticity which is essential for fundamental processes like cognition, memory, and learning(*3*). Members of the Shank gene family function as master scaffolding proteins in the post-synaptic density of excitatory neurons assembling the PSD-95 protein (postsynaptic density-95), NMDA receptors, AMPA receptors, and homer-based complexes(*2*, *4*). Specifically, the impairment of *Shank3* function has been primarily associated with the Phelan-McDermid syndrome (PMS)(*5*), also known as the 22q13.3 deletion syndrome. PMS is a neurodevelopmental condition caused by a deletion of the region spanning the *Shank3* gene (the distal arm of Chromosome 22)(*6*) and characterized by delayed speech and language development, ASD-like symptoms, intellectual disabilities (ID), and other behavioral abnormalities(*6*, *7*). Some PMS patients exhibit symptoms identical to those reported in patients with haploinsufficiency of *Shank3* mutations, implying a correlation between *Shank3* function loss and the manifestation of specific neurodevelopmental symptoms in PMS patients(*7*).

Apart from PMS, several studies have also revealed point mutations in *Shank3* in patients with ASDs, IDs, and Schizophrenia (SCZ) (*8–10*). Mutations in the *Shank3* gene are strongly associated with ASD with estimates of approximately 1% of the incidence in ASD patients(*11*). Moreover, transgenic mice models with a *Shank3* knockout or loss of function exhibit ASD symptoms such as decreased social interactions, repetitive behaviors, cognitive deficits, and synaptic dysfunction(*12*, *13*). We recently generated iPSCs from a PMS patient with a *Shank3* mutation (c.3679insG)(*14*) and reported the neurophysiological properties of iPSC-derived cortical neurons(*15*). Our results demonstrated that cortical neurons with this *Shank3* mutation, similar to several other ASD-associated mutations, displayed early maturation and hyperexcitability in comparison to unaffected healthy neurons. Neuronal hyperexcitability due to *Shank3* haploinsufficiency was also reported by another group in human embryonic stem cell-derived glutamatergic neurons as well as in mouse primary hippocampal neurons harboring *Shank3* deletion(*16*). Hyperactivity of the corticostriatal circuit and abnormal maturation were also observed during the early development of *Shank3B*^-^/^-^ mice displaying autistic characteristics(*17*).

In recent years, EVs and their role in neurological disorders have emerged as a novel area of research. EVs are membrane-bound, nano-sized vesicles released primarily by eukaryotic cell types, including neural cells(*18*). EVs transport a wide range of proteins, lipids, and nucleic acids from the parent cells, making them useful intercellular communication players under both healthy and diseased conditions(*19*). Several reports have emphasized the role of EVs as mediators of signaling molecules during immunological and inflammatory cellular responses(*20*). EVs have also been reported to transport misfolded or aggregated proteins associated with neurodegenerative diseases such as Alzheimer’s disease (AD), Parkinson’s disease (PD), and amyotrophic lateral sclerosis (ALS) to the recipient cells leading to the dissemination of pathological features(*21*). Previous research has also documented the beneficial properties of EVs by demonstrating that EVs originating from oligodendrocytes enhance neuronal survival under conditions of cellular stress(*22*, *23*). EVs derived from stem cells, notably mesenchymal stem cells (MSCs) and iPSCs have garnered significant interest due to their therapeutic potential(*24*, *25*). Unlike stem cell transplantation therapies, EVs derived from these cells are generally non-immunogenic, can be administered via multiple routes (e.g. intranasal, intravenous, and using a nebulizer), and pose minimal ethical concerns(*26*). However, despite ample evidence, the therapeutic application of EVs in neurological disorders remains largely constrained, primarily due to the lack of preclinical studies conducted using human cells(*27*). Moreover, the mechanisms by which EVs impart their neuroprotective effect are underexplored.

In this study, we utilized an iPSC model of the autism-associated c.3679insG *Shank3* mutation and a transgenic *Shank3B-/-* mouse model to explore the impact of EVs on cellular neurophysiological properties and mice behavioral phenotypes. We hypothesized that the early hyperexcitability observed in *Shank3* mutant neurons might be mediated by extracellular vesicles, which we tested in an EV exchange experiment. We discovered that when treating control neurons with EVs derived from *Shank3* mutant neurons, they exhibited properties like *Shank3* mutant neurons with an increased excitability while *Shank3* mutant neurons treated with EVs derived from control neurons did not change and remained hyperexcitable. In addition, to explore the therapeutic potential of stem cell-derived EVs, we treated *Shank3* mutant cortical neurons with EVs derived from MSCs (MSC-EVs) and iPSCs (iPSC-EVs) both from healthy control donors. We found that both MSC-EVs and iPSC-EVs were able to rescue the abnormal electrophysiological properties of *Shank3* mutant cortical neurons, making them similar to control neurons. Finally, as a further step towards possible clinical use, we treated *Shank3B-/-* mice by intranasal administration of iPSC-derived EVs from early postnatal to the juvenile stage. This treatment rescued ASD-associated behavioral deficits in the *Shank3B-/-* mice. Proteomic analysis revealed distinct EV cargoes especially synaptic and plasticity regulators that could be mediating these neurophysiological effects in respective recipient neurons. Overall, our study highlights iPSC-derived EVs as modulators of neuronal physiological properties and a potential therapeutic candidate for ASD.

## RESULTS

### Characterization of EVs from human cortical neurons derived from control and *Shank3* (c3679insG) iPSC lines

*Shank3* is ∼190 kDa protein with a multi-domain architecture including an SH3 domain, a proline-rich domain, a PDZ domain, and ankyrin repeat domains^35^. The c3679insG heterozygous mutation of *Shank3* previously reported to cause ASD is located in the Exon-21 of *Shank3*(*28*) (Proline-rich domain) and was further characterized in the transgenic mice model (*13*)(Fig 1a,b). Single-cell RNA sequencing data available from the UCSC browser reveals that *Shank3* is abundantly expressed in the cortex and cerebellum regions of the human fetal & adult brain; *Shank3* is comparatively expressed more in excitatory and subtypes of inhibitory neurons than astrocytes, oligodendrocytes, and microglial cells (Fig 1c).

**Figure 1.**
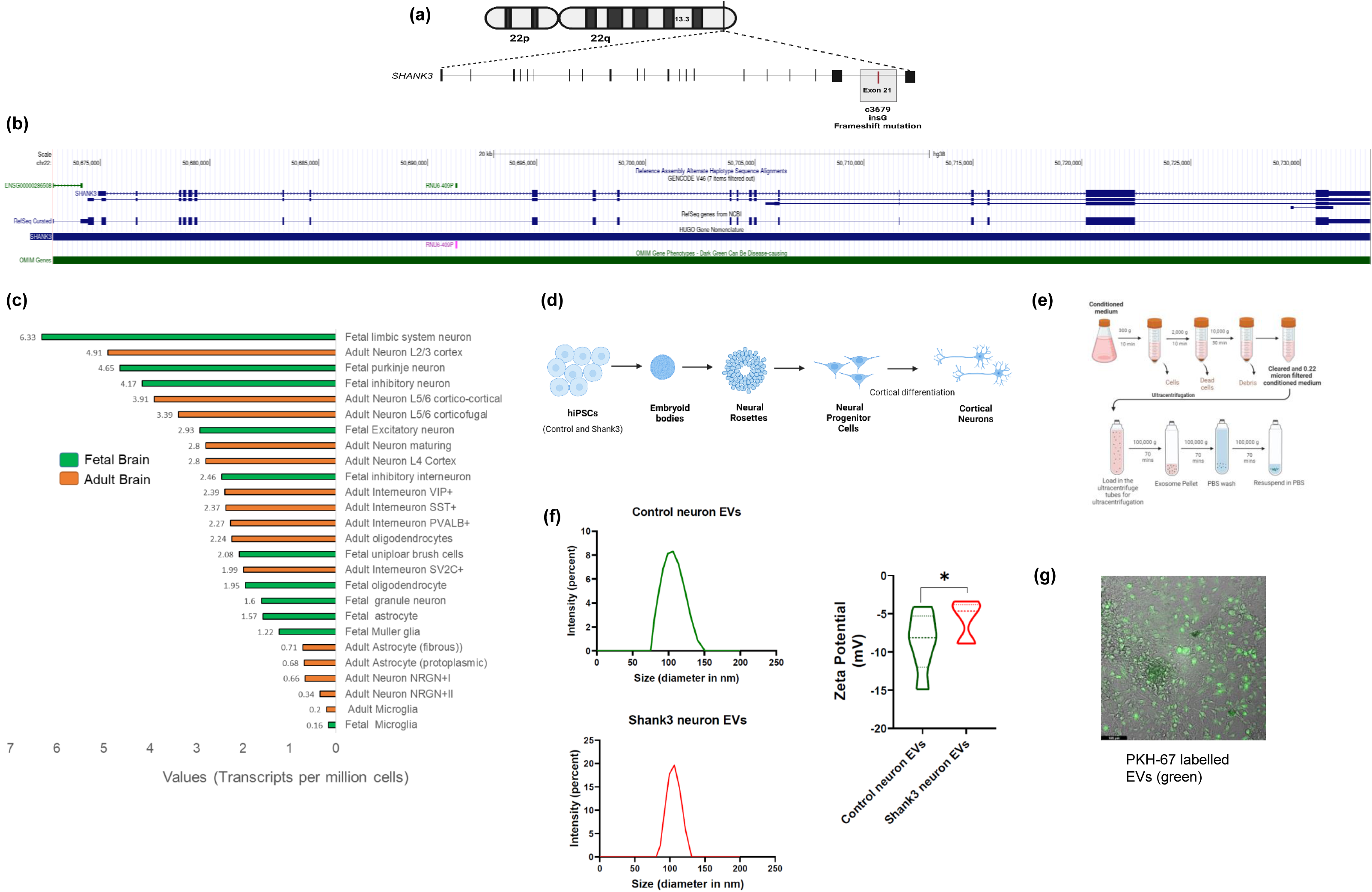
Characterization of EVs isolated from cortical neurons derived from control and *Shank3* (c3679insG) iPSC lines. (a) An illustration of a frameshift mutation in the *Shank3* (c3679insG) gene causing PMS and ASD symptoms in a female child(*14*). (b) A snapshot of the UCSC browser(*69*) for the human *Shank3* gene coordinates at Chromosome 22 and NCBI ref-seq gene architecture using hg38 genome assembly. (c) The expression of *Shank3* derived from single-cell RNA sequencing experiments plotted for the fetal and adult human brain. The data was obtained from the single-cell RNA expression track settings (publicly available, last updated on 2022-04-11) from the UCSC browser(*69*). (d) Schematics for cortical differentiation from control and *Shank3* mutant human iPSCs by differentiation into embryoid bodies (EBs), neural progenitor cells (NPCs), and then into cortical neurons. The cortical differentiation from NPCs was performed for 29-31 days. (e) The extracellular vesicle isolation procedure used in our study included differential centrifugation to remove dead cells and debris from the conditioned media from control and *Shank3* mutant neurons followed by ultracentrifugation (see methods). (f) Size and zeta potential quantification by Zetasizer^TM^ using dynamic light scattering of Control and *Shank3* neuron-derived EVs. (g) A micrograph for uptake of PKH-67 labeled EVs by cortical neurons derived from control iPSC lines.

We recently generated iPSC lines with *Shank3* c3679insG mutation and control from a first-degree relative(*14*). We further differentiated these iPSC lines into cortical NPCs and neurons (Fig 1d for the schematics) (see methods). We isolated EVs from control and *Shank3* cortical neurons (see methods) and characterized the size and zeta potential by dynamic light scattering (Fig 1e,f). The size of EVs derived from control neurons was 108±20.6 nm and EVs derived from *Shank3* mutant neurons were 103±13.5 nm (p=0.63). EVs derived from control neurons had an average zeta potential of −8.6±2.4 mV while the EVs derived from *Shank3* mutant neurons had an average zeta potential of −5.7±0.45 mV (p=0.039). We labeled the isolated EVs using the PKH-67 dye and verified the uptake of EVs in control cortical neurons (Fig 1g).

According to the Minimal Information for Studies of Extracellular Vesicles (*MISEV*) 2023 guidelines(*29*), protein characterization of EVs from control and *Shank3* mutant neurons revealed distinct marker profiles (Category 1, 2, and Category 5) consistent with *MISEV* recommendations (Supplementary Table 1) and the absence of non-EV markers of category 3 (ER proteins like *Calnexin*, mitochondrial proteins *VDAC*, *Cytochrome C* or Golgi bodies protein like *GM130*).

### EVs derived from *Shank3* mutant neurons induce *Shank3* phenotypes in control neurons

As previously reported, cortical neurons derived from *Shank3* mutant iPSCs exhibited an accelerated maturation with early hyper-excitability, increased Na+ currents, and increased rate of excitatory postsynaptic currents (EPSCs) compared to control neurons when the neurons were differentiated for 4-5 weeks(*15*). We performed the switching of EVs two times between control and *Shank3* mutant neurons during the differentiation (see methods) and measured the effect on the electrophysiological characteristics of the cells (Fig 2a, schematic of the experiment); i.e., the control cortical neurons were treated with EVs derived from *Shank3* mutant neurons and vice-versa. We further performed immunostaining with neuronal marker *MAP2* and cortical neuronal markers *TBR1* and *CTIP2* for untreated and extracellular vesicle-treated control neurons (Fig 2b). ICC shows 35±5% *TBR1*+ neurons and 9±1.5% *CTIP2*+ neurons in the differentiated neuronal cultures (among *MAP2*+ cells) in untreated control neurons and 47±11% *TBR1*+ and 11±2% *CTIP2*+ (among *MAP2*+ cells) in control neurons treated with EVs derived from *Shank3* mutant neurons (Supplementary Fig 1).

**Figure 2.**
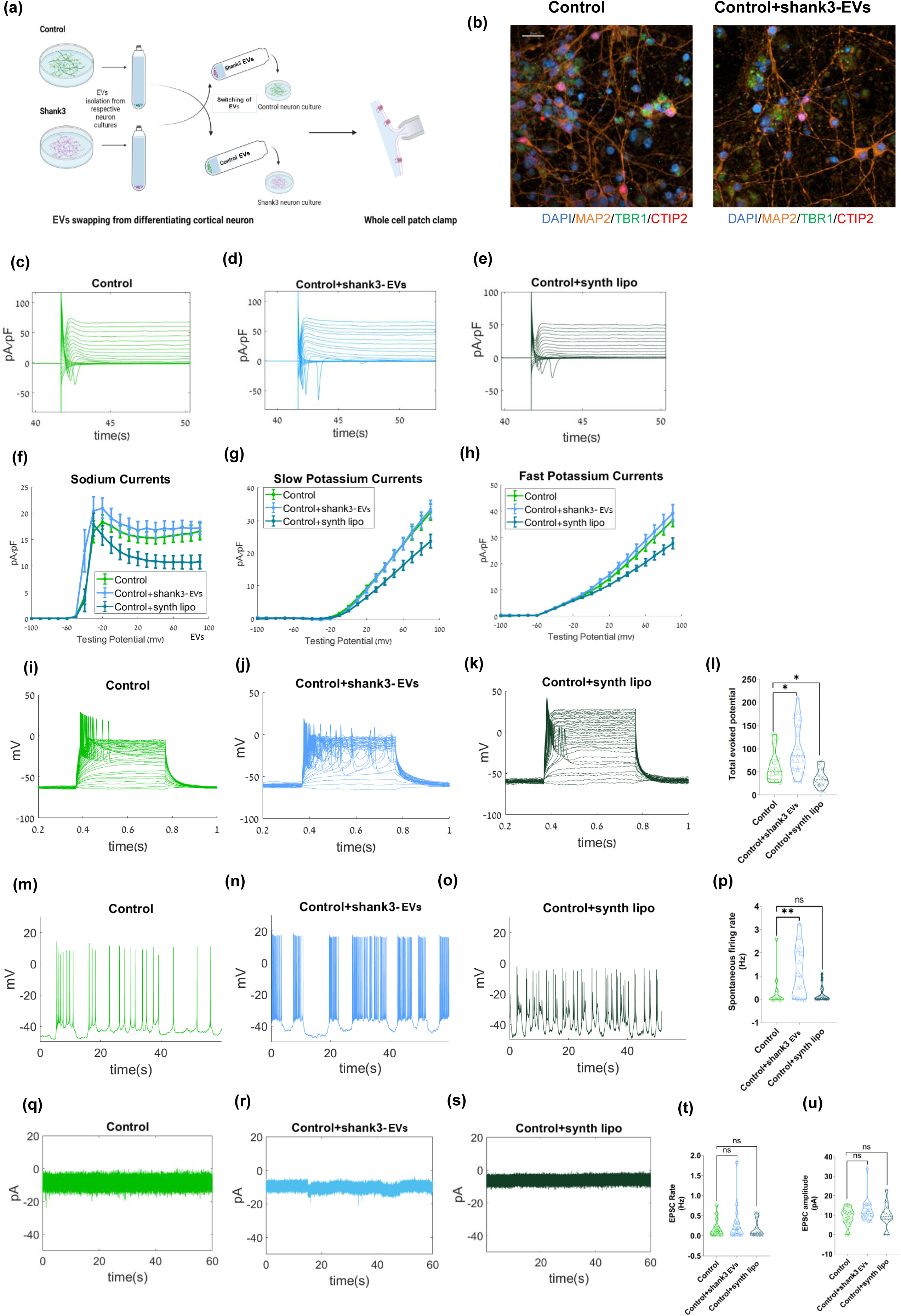
EVs derived from *Shank3* mutant neurons induce *Shank3* phenotypes in control cortical neurons. (a) A schematic of the experiment: EVs were isolated from control and *Shank3* mutant cortical neurons. The purified EVs were exchanged. i.e., control neurons were treated with EVs derived from *Shank3* mutant neurons and vice-versa. The functional analysis of neurons was performed using a whole-cell patch-clamp after ∼30 days of differentiation. (b) Immunocytochemistry images of cortical markers *TBR1* (green), *CTIP2* (red); panneuronal marker *MAP2* (orange), and DAPI (blue) for (left) control neurons and (right) control neurons treated with EVs derived from *Shank3* mutant neurons. *See supplementary* figure 1 *for quantification*. The EVs were added at two time points (see methods); Scale bar, 20 µM. (c-e) Na+/K+ currents example traces recorded in voltage clamp mode in (c) control cortical neurons (d) control cortical neurons treated with EVs derived from *Shank3* mutant neurons, and (e) control cortical neurons treated with synthetic liposomes. (f) The average Na+ current recorded in the control neurons treated with EVs derived from *Shank3* mutant neurons (light blue) showed an increasing trend while control neurons treated with synthetic liposomes (dark green) were significantly lower compared to control neurons (green). (g) Similarly, the average slow K+ currents, and (h) the average fast K+ currents. (i-k) Evoked potentials example traces are shown in (i) control cortical neurons, (j) control cortical neurons treated with EVs derived from *Shank3* mutant neurons, and (k) control cortical neurons treated with synthetic liposomes. (l) A violin plot of the total evoked potentials for control neurons treated with EVs derived from *Shank3* mutant neurons (light blue) showed increased APs while control neurons treated with synthetic liposomes (dark green) had fewer APs compared to control neurons (green). (m-o) Spontaneous firing example traces are shown in (m) control neurons, (n) control neurons treated with EVs derived from *Shank3* mutant neurons, and (o) control cortical neurons treated with synthetic liposomes. (p) A violin plot for the average spontaneous firing rate in control neurons treated with EVs derived from *Shank3* mutant neuron*s* (light blue) shows an increased firing rate compared to control neurons (green), and control neurons treated with synthetic liposomes (dark green). (q-s) EPSCs example traces are shown in (q) control neurons, (r) control neurons treated with EVs derived from *Shank3* mutant neurons, and (s) control neurons treated with synthetic liposomes. (t-u). The violin plots for the average EPSCs rate (t), and EPSCs amplitude (u) measured in control neurons (green), control neurons treated with EVs derived from *Shank3* mutant neuron*s* (light blue), and control neurons treated with synthetic liposomes (dark green) show no significant changes; \**p*<0.05, \*\**p*<0.01.

After EV switching, we performed electrophysiological recordings when the neurons were in the fifth week of *in-vitro* differentiation in EV-treated and untreated control neurons (Fig 2). The control cortical neurons displayed normal Na+/K+ currents of young, immature cortical neurons (Fig 2c, shows a representative trace). Comparatively, although not statistically significant control neurons treated with EVs derived from *Shank3* mutant neurons showed a trend toward increased Na+ currents after EV swapping (p=0.49) (Fig 2d,f). To determine whether the observed effects were specific to EV treatment and not simply due to exposure to nano-sized lipid membrane particles, we included an additional treatment using synthetic liposomes (∼100 nm in size; see methods) in control cortical neurons (Fig 2e shows a representative trace and Fig 2f shows the averages). Interestingly, synthetic liposomes elicited the opposite effect of treatment with EVs derived from *Shank3* mutant neurons, resulting in a significant reduction of Na+/K+ currents in control cortical neurons (p= 0.0005, Na+; p=0.000003, slow K+; p=0.00001, fast K+) (Fig 2f–h).

We then measured the evoked action potentials (APs) in the current clamp mode. The representative traces of evoked APs for control cortical neurons and control neurons treated with EVs derived from *Shank3* mutant neurons and synthetic liposomes are shown in Fig 2i-k. Surprisingly, control neurons treated with EVs derived from *Shank3* mutant neurons displayed an increased number of evoked APs and were hyper-excitable compared to the control cortical neurons (Fig 2l presents the averages). The synthetic liposome treatment, however, caused a decrease in the number of evoked APs and hence the excitability (Fig 2l). Next, we investigated the spontaneous firing rate of neurons when holding the cell at −45 mV (which is approximately the resting membrane potential of the immature neurons). The representative traces of the spontaneous APs in control neurons, control neurons treated with EVs derived from *Shank3* mutant neurons, and synthetic liposomes-treated control neurons have been plotted in Fig 2m-o. The rate of spontaneous APs was higher in control neurons treated with EVs derived from *Shank3* mutant neurons compared to the untreated control neurons (Fig 2p presents the averages and Fig 2n presents a representative example). The synthetic liposome treatment decreased the rate of spontaneous APs (not significantly) and hence had a contrasting effect compared to EVs derived from *Shank3* mutant neurons (Fig 2o,p).

We further measured EPSC events in voltage-clamp mode (see methods) to analyze the synaptic activity of control neurons at the fifth week of cortical differentiation. The representative traces of EPSC rate and amplitudes for untreated control neurons, control neurons treated with EVs derived from *Shank3* mutant neurons, and synthetic liposomes treated control neurons have been plotted in Fig 2q-s. Overall, at this stage of the differentiation, the EPSC rate and amplitude were not significantly changed by any of these treatments (Fig 2t,u shows the averages).

### EVs derived from control neurons do not rescue the phenotype of *Shank3* cortical neurons

We similarly measured and analyzed the electrophysiological properties of *Shank3* mutant neurons after EV switching. The immunostaining performed in *Shank3* mutant neurons and *Shank3* mutant neurons treated with EVs derived from control neurons showed (48±3% *TBR1*+/*MAP2*+; 23±3% *CTIP2*+/*MAP2*+) and (38±8% *TBR1*+/*MAP2*+; 14±4% *CTIP2*+/*MAP2*+) respectively (Fig 3a & Supplementary Fig 2a,3b). *Shank3* mutant neurons treated with EVs derived from control neurons had similar Na+/K+ currents to untreated *Shank3* mutant neurons (Fig 3b-f). We have previously shown that *Shank3* mutant neurons display an increased amplitude of normalized Na+ currents(*15*). This was also evident in this study’s patch clamp recordings (Fig 3g-i). The number of evoked action potentials (APs) measured in current clamp mode in *Shank3* mutant neurons showed an increased number of spikes when compared to control neurons (p=0.03; averages for combined APs data from previous study(*15*)) (Fig 3m). However, treatment with EVs derived from control neurons did not reduce the hyperexcitability of *Shank3* mutant neurons (Fig 3j-l).

**Figure 3.**
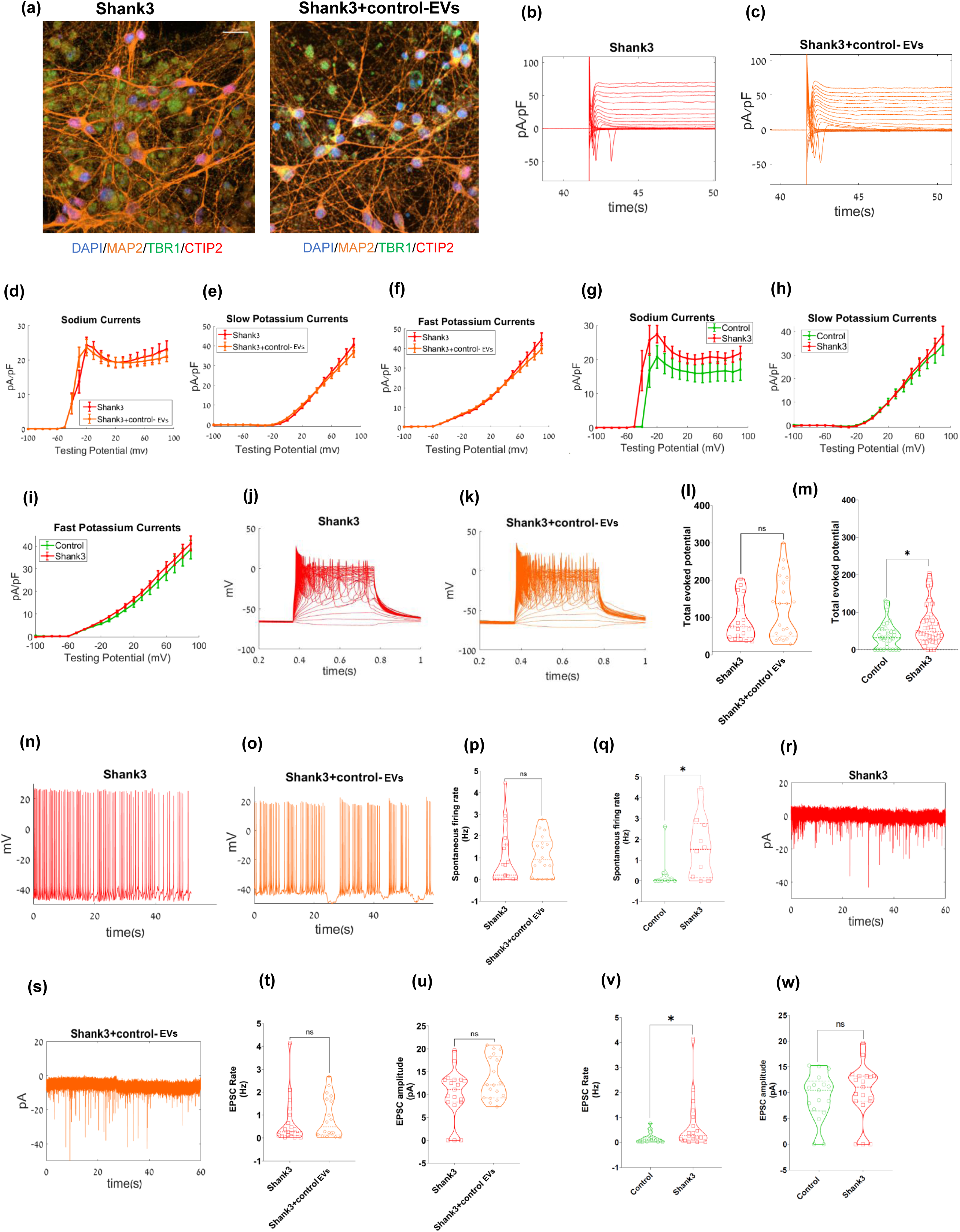
EVs derived from control neurons do not rescue the phenotypes of *Shank3* mutant cortical neurons. (a) Immunocytochemistry images of *TBR1* (green), *CTIP2* (red), *MAP2* (orange), and DAPI (blue) for untreated *Shank3* mutant neurons (left) and *Shank3* mutant neurons treated with EVs derived from control neurons (right). *See supplementary* figure 1 *for quantification*. The EVs were added at two time points (see methods); Scale bar, 20 µM. (b-c) Na+/K+ currents example traces are shown for (b) *Shank3* mutant neurons and (c) *Shank3* mutant cortical neurons treated with EVs derived from control neurons after ∼30 days of differentiation. (d) The average Na+ currents recorded in *Shank3* mutant neurons (red) and *Shank3* mutant neurons that were treated with EVs derived from control neurons (orange) did not change significantly. (e) Similarly, the average slow K+ currents, and (f) The average fast K+ currents. (g-i) The average Na+ currents (g) recorded in the *Shank3* mutant neurons (red) were significantly higher compared to control neurons (green). Similarly, the averages of the (h) slow K+ currents and (i) fast K+ currents are presented. Evoked potentials example traces are presented for (j) *Shank3* mutant neurons (red) and (k) *Shank3* mutant neurons treated with EVs derived from control neurons (orange). (l) A violin plot of the average total evoked potentials for *Shank3* mutant neurons (red) and *Shank3* mutant neurons treated with EVs derived from control neurons (orange) shows no significant change. (m) A violin plot showing significantly higher total evoked APs for *Shank3* mutant neurons (red) compared to control neurons (green) after ∼30 days of differentiation. The APs data from the previous study(*15*) was also combined and analyzed together (control, n=32; *Shank3* mutant, n=34). Spontaneous firing example traces are shown for (n) *Shank3* mutant neurons (red) and (o) *Shank3* mutant neurons treated with EVs derived from control neurons (orange). (p) A violin plot for the average spontaneous firing rate for *Shank3* mutant neurons (red) and *Shank3* mutant neurons treated with EVs derived from control neurons (orange) with no significant change. (q) Similarly, a violin plot for the average spontaneous firing rate for *Shank3* mutant neurons (red) with a higher firing rate compared to control neurons (green). EPSC example traces can be observed for (r) *Shank3* mutant neurons (red), and (s) *Shank3* mutant neurons treated with EVs derived from control neurons(orange). A violin plot for the (t) EPSC rate, and (u) EPSC amplitude for *Shank3* mutant neurons (red), and *Shank3* mutant neurons treated with EVs derived from control neurons (orange) are presented without any significant change. Similarly, violin plots for the *Shank3* mutant neurons (red) (v) with significantly higher EPSC rate and (w) no change in EPSC amplitude compared to control neurons (green) after ∼30 days of differentiation are shown; \**p*<0.05, \*\**p*<0.01. NS: not significant.

Similarly, we recorded spontaneous APs by holding the cells at −45mV. The average rate of spontaneous APs was significantly higher in *Shank3* mutant neurons compared to control neurons (Fig 3q). Notably, EVs derived from control neurons did not reduce the spontaneous APs firing rate of the *Shank3* mutant neurons (Fig 3n-p). The EPSC rate was also significantly higher in *Shank3* mutant neurons compared to the control neurons (p=0.04) (Fig 3v,w; averages of EPSC rate and amplitude) as previously reported(*15*). *Shank3* mutant neurons maintained a high EPSC rate and amplitude after treatment with EVs derived from control neurons (Fig 3r- u shows representative traces and averages).

Thus, these results (Fig 2 and 3 collectively) show that treatment with EVs derived from *Shank3* mutant neurons can significantly influence the neurophysiology of control cortical neurons and make them hyper-excitable, whereas treatment with EVs derived from control neurons does not have a significant effect on *Shank3* mutant neurons’ physiology.

### Mesenchymal stem-cell-derived EVs rescue *Shank3* cortical neuronal phenotypes

Mesenchymal stem cell (MSC) derived EVs (MSC-EVs) have been previously shown to have rescued behavioral deficits in mouse models of Autism(*30*) and Schizophrenia(*31*). Hence, we wanted to examine if MSC-EVs can rescue the neurophysiological deficits of human *Shank3* mutant cortical neurons. We isolated EVs from MSCs (see methods) and treated *Shank3* mutant neurons (see methods) (Fig 4a, shows the schematics of the experiment) with purified MSC-EVs (∼ 10^5^ EV particles/cell), and measured the electrophysiological properties of the treated neurons using whole cell patch clamp. The average size of MSC-EVs was 73.4±12.5 nm (Fig 4b shows the size distribution) and the zeta potential was -16.7±1.3 mV. MSC-EVs were characterized by canonical EV markers *B2M*, *CD63*, *CD81*, *CHMP4A*, and *PDCD6IP*, along with the MSC-specific marker *NT5E* (Supplementary Table 1). Immunostaining was performed for *MAP2*, *TBR1*, and *CTIP2* in the untreated and MSC-EV-treated *Shank3* mutant neurons (Fig 4c and Supplementary Fig 2b). The expression was measured to be 31±5% *TBR1*+/*MAP2*+; 11±2% *CTIP2*+/*MAP2*+ in MSC-EVs treated *Shank3* mutant neurons and 48±3% *TBR1*+/*MAP2*+; 23±3% *CTIP2*+/*MAP2*+ in untreated *Shank3* mutant neurons (Supplementary Fig 2b,3b).

**Figure 4.**
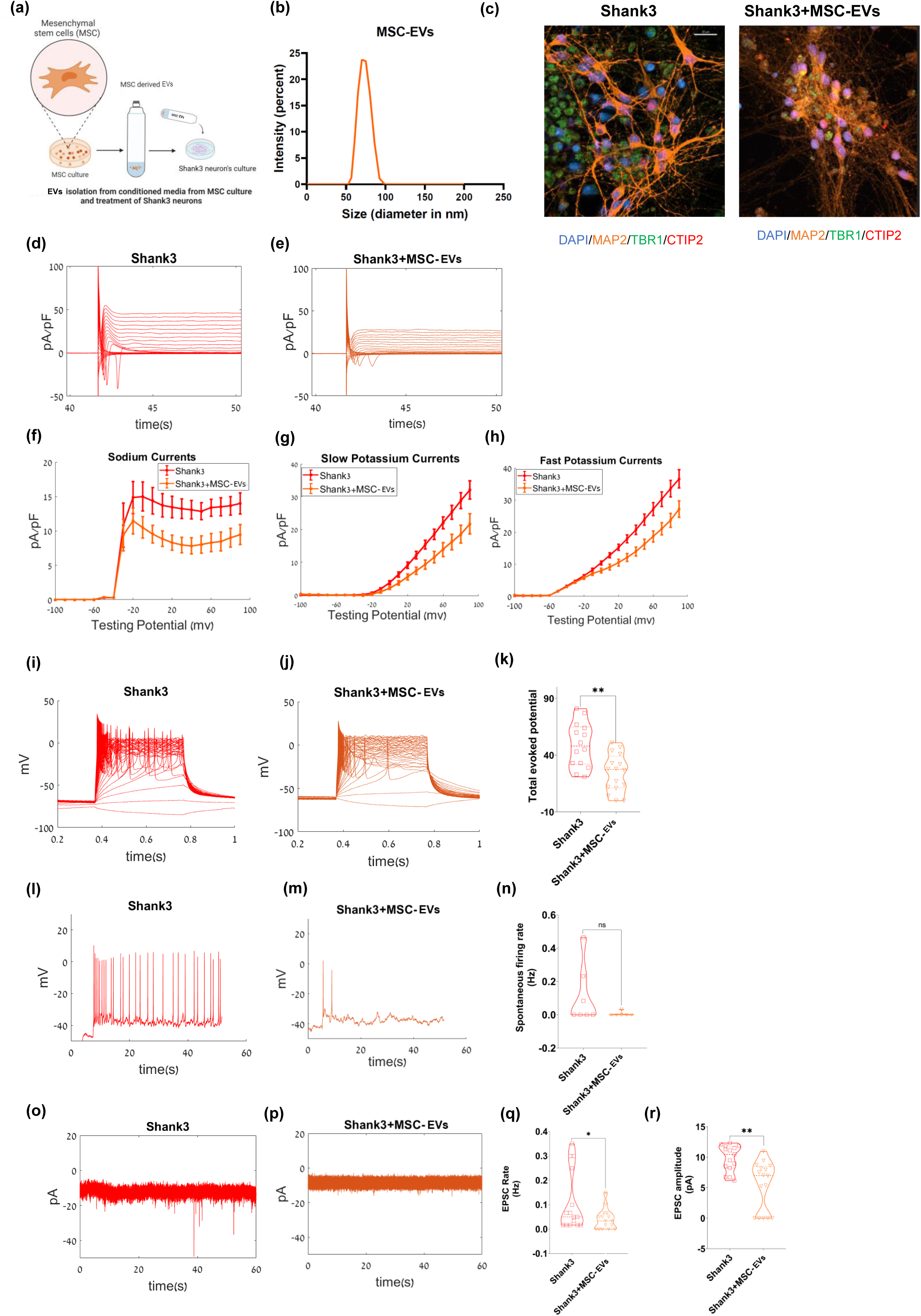
Mesenchymal stem-cell-derived EVs rescue *Shank3* cortical neuronal phenotypes. (a) A schematic demonstrating the isolation of EVs from the conditioned media of mesenchymal stem cell (MSC) culture and treatment of the *Shank3* mutant cortical neurons with MSC-EVs (see methods). The treatment with MSC-EVs was performed three times during the cortical differentiation timeline. The electrophysiological parameters were measured using the whole cell patch clamp technique at ∼30 days of differentiation. (b) Size characterization of MSC-EV by dynamic light scattering. (c) Immunocytochemistry images of *TBR1* (green), *CTIP2* (red), *MAP2* (orange), and DAPI (blue) of untreated *Shank3* mutant neurons (left) and *Shank3* mutant neurons treated with MSC-derived EVs (right); Scale bar, 20 µM. Na+/K+ currents example traces are shown in (d) *Shank3* mutant neurons and (e) *Shank3* mutant neurons treated with MSC-derived EVs. (f) The average Na+ currents were recorded in *Shank3* mutant neurons (red) and *Shank3* treated with MSC-derived EVs (orange). Similarly, (g) the average slow K+ and (h) the average fast K+ currents are presented. Example traces for evoked potentials are shown in (i) *Shank3* mutant neurons (red) and (j) *Shank3* mutant neurons treated with MSC-derived EVs (orange). (k) A violin plot for the total evoked potentials with a significant decrease in APs for *Shank3* mutant neurons treated with MSC-derived EVs (orange) compared to *Shank3* mutant neurons (red). Example traces for spontaneous firing are shown in (l) *Shank3* mutant neurons (red) and (m) *Shank3* mutant neurons treated with MSC-derived EVs. (n) A violin plot for the average spontaneous firing rates for *Shank3* mutant neurons (red) and *Shank3* mutant neurons treated with MSC- derived EVs (orange) showed no significant change. EPSC example traces can be observed in (o) *Shank3* mutant neurons (red), and (p) *Shank3* mutant neurons treated with MSC-derived EVs (orange). A violin plot showed a significant decrease for the averages of (q) EPSC rate and (r) EPSC amplitude in *Shank3* mutant neurons treated with MSC-derived EVs (orange) compared to *Shank3* mutant neurons (red) after ∼30 days of differentiation are presented; \**p*<0.05, \*\**p*<0.01. NS: not significant.

Fig 4d-e shows representative traces for the normalized Na+/K+ currents (in voltage-clamp mode) of *Shank3* mutant cortical neurons with and without MSC-EV treatment at 4-5 weeks of *in-vitro* differentiation. MSC-EVs treatment of *Shank3* mutant neurons significantly reduced the Na+ by ∼30% (p=2.37e-09), fast K+ currents by ∼30% (p= 0.0001), and slow potassium by ∼30% (p=0.0004) (Fig 4f-h). The number of evoked APs (measured in current-clamp mode) also indicated a ∼ 40% reduction in the excitability of *Shank3* mutant neurons (p=0.002) (Fig 4i-k for representative traces and averages) compared to untreated mutant neurons. Additionally, the rate of spontaneous APs (see methods) showed a decreasing trend but a non-significant change (p=0.29) after MSC-EVs treatment (Fig 4l-n for representative images and averages).

We then analyzed EPSCs in voltage-clamp mode and found a significant reduction in the EPSC rate (p=0.049) and amplitude (p=0.0009) in MSC-EV-treated *Shank3* mutant neurons compared to untreated *Shank3* cortical neurons (Fig 4o-r for representative traces and averages). These results show that EVs isolated from MSCs rescue most *Shank3* mutant cortical neuronal phenotypes.

### iPSC-derived EVs rescue *Shank3* cortical neuronal phenotypes

Subsequently, we aimed to examine if EVs derived from control iPSCs (iPSC-EVs) could also rescue the phenotypes of *Shank3* mutant cortical neurons compared to untreated neurons (Fig 5a shows the schematics of the experiment). Hence, we isolated EVs from the iPSC culture medium and treated the *Shank3* mutant neurons with ∼10^5^ EV particles/cell (see methods). Then, we measured the neurophysiological properties using a whole cell patch clamp. The average size of the isolated EVs was 70.3±7.8 nm and the zeta potential was -20.7±1.02 respectively (Fig 5b). iPSC-derived EVs contained *ANXA1*, *GAPDH*, and *FGF2*, alongside epithelial markers *DSG1*, and *DSC1* as per *MISEV 2023* guidelines(*29*) (Supplementary Table 1). Fig 5c shows immunostaining for untreated *Shank3* mutant neurons and iPSC-EV-treated *Shank3* mutant neurons. The quantification of different cortical markers showed 48±3% *TBR1*+/*MAP2*+; 23±3% *CTIP2*+/*MAP2*+ in untreated *Shank3* mutant neurons and 34±3% *TBR1*+/*MAP2*+; 10±1% *CTIP2*+/*MAP2*+ in iPSC-EVs treated *Shank3* mutant neurons (also see Supplementary Fig 3a,3b for the quantification).

**Figure 5.**
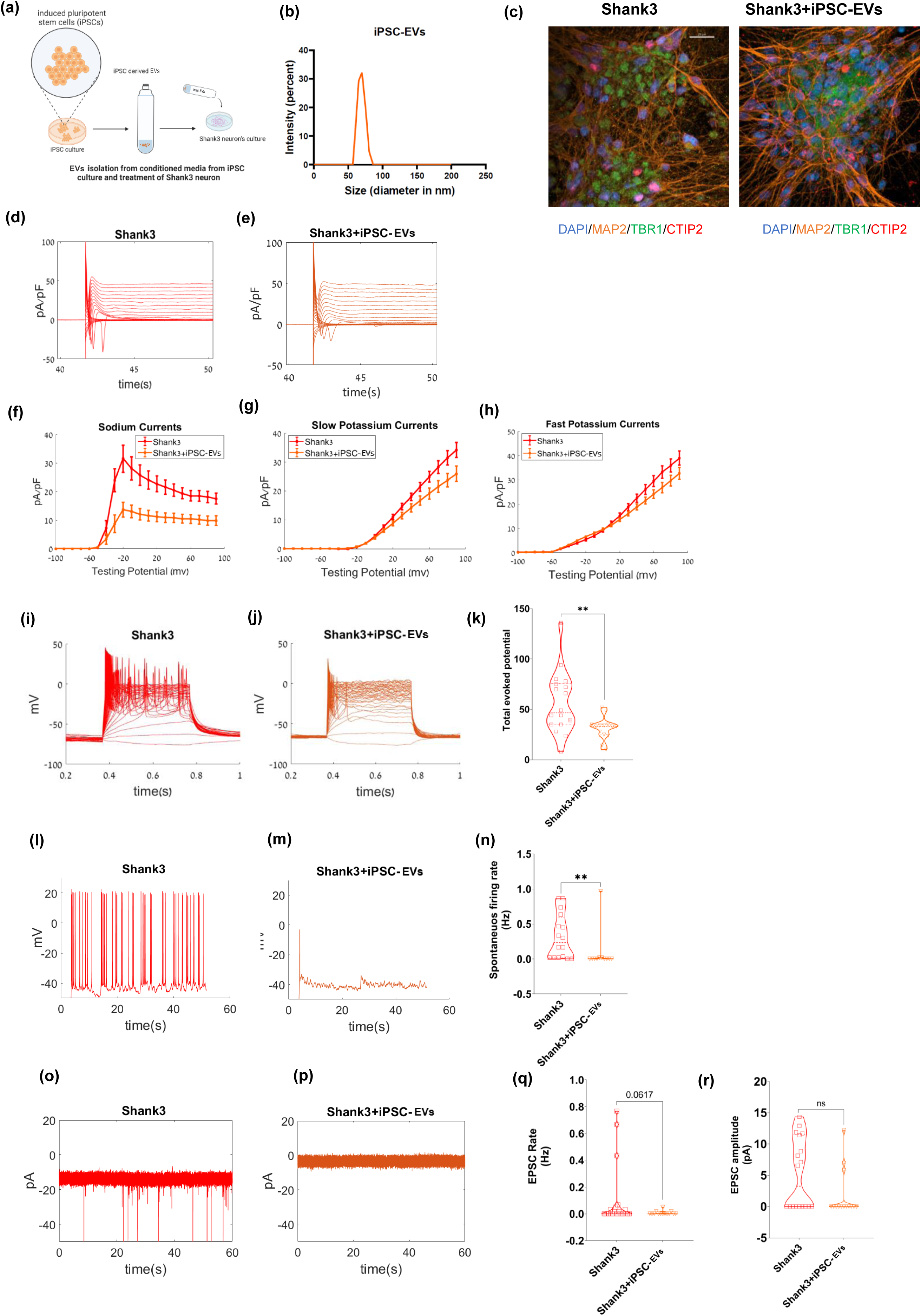
iPSC-derived EVs rescue *Shank3* cortical neuronal phenotypes. (a) A schematic demonstrating the isolation of EVs from the conditioned media of the induced pluripotent stem cell (iPSC) culture and treatment of *Shank3* mutant cortical neurons with iPSC-EVs during the differentiation. The treatment with iPSC-EVs was performed three times during the cortical differentiation timeline. The electrophysiological parameters were measured using the whole cell patch clamp technique at ∼30 days of differentiation. (b) Size characterization of iPSC-EVs by dynamic light scattering. (c) Immunocytochemistry images of *TBR1* (green), *CTIP2* (red), *MAP2* (orange), and DAPI (blue) for untreated *Shank3* mutant neurons (left) and *Shank3* mutant neurons treated with iPSC-derived EVs (right); Scale bar, 20 µM. Na+/K+ currents example traces are shown for (d) *Shank3* mutant cortical neurons and (e) *Shank3* mutant neurons treated with iPSC-derived EVs. (f) The average Na+ currents recorded in *Shank3* mutant neurons treated with iPSC-derived EVs (orange) were significantly reduced compared to *Shank3* mutant neurons (red). Similarly, (g) the average slow K+ and (h) the average fast K+ currents were also reduced. Example traces for evoked potential recorded in (i) *Shank3* mutant neurons (red) and (j) *Shank3* mutant neurons treated with iPSC-derived EVs (orange) are presented. (k) A violin plot for the total evoked potentials for *Shank3* mutant neurons treated with iPSC- derived EVs (orange) showed a reduced number of APs compared to *Shank3* mutant neurons (red) Example traces for spontaneous firing recordings are presented for (l) *Shank3* mutant neurons (red) and (m) *Shank3* mutant neurons treated with iPSC-derived EVs. (n) A violin plot for the average spontaneous firing rate for *Shank3* mutant neurons treated with iPSC-derived EVs (orange) showed a significantly lower firing rate compared to *Shank3* mutant neurons (red). EPSC example traces can be observed for (o) *Shank3* mutant neurons (red) and (p) *Shank3* mutant neurons treated with iPSC-derived EVs (orange). A violin plot for (q) EPSC rate and (r) EPSC amplitude in *Shank3* mutant neurons (red) and *Shank3* mutant neurons treated with iPSC-derived EVs (orange) showed no significant change after ∼30 days of differentiation; \**p*<0.05, \*\**p*<0.01. NS: not significant.

The normalized Na+ and K+ currents of *Shank3* mutant neurons were significantly reduced after iPSC-EV treatment compared to untreated neurons measured in voltage-clamp mode as shown in representative traces and average plots (Na+, p=3.09e-08; fast K+, p= 0.008; slow K+, p=0.002) (Fig 5d-h). The total number of evoked APs (measured in current clamp mode, see methods) also demonstrated a reduction in the hyper-excitability phenotype in *Shank3* mutant neurons after iPSC-EV treatment (p=0.004) as shown in representative traces and average plots (Fig 5i-k). Similarly, the rate of spontaneous APs (see methods) was decreased in the iPSC-EV treated neurons compared to untreated *Shank3* mutant neurons (p=0.0028) (representative and average plots in Fig 5l-n).

Finally, we measured the EPSC rate and amplitude (in voltage clamp mode, see methods) in the iPSC-EV-treated *Shank3* mutant neurons compared to untreated *Shank3* mutant neurons. These were reduced but not significantly (EPSC rate, p=0.06; EPSC amplitude, p=0.07) (Fig 5o-r). Overall, our results show that EVs derived from iPSCs rescue most of the neurophysiological phenotypes in *Shank3* mutant neurons making them more similar to control neurons.

### Proteomic analysis of functional cargoes of extracellular vesicles

To better understand the molecular mechanisms that may be involved in the alterations in neurophysiology, we performed quantitative proteomics using LC-MS/MS to identify the protein cargoes of the four types of EVs that we isolated for the treatment of control and Shank3 mutant neurons. These included EVs derived from control neurons, EVs derived from Shank3 mutant neurons, MSC-EVs, and iPSC-EVs. We detected a larger number of proteins in EVs derived from *Shank3* mutant neurons (216 proteins) and MSC–EVs (217 proteins) as compared to EVs derived from control neurons (114 proteins) and iPSC-EVs (32 proteins) (Fig 6a presents a Venn diagram). A cellular component analysis of protein cargoes from all four cell sources showed significant enrichment for “extracellular vesicles” and “extracellular “ proteins (p<0.001, Fig 6b). Additionally, the neuron-derived EVs also showed enrichment for “cytoplasm” (p<0.001), but not the stem cell-derived extracellular vesicles.

**Figure 6.**
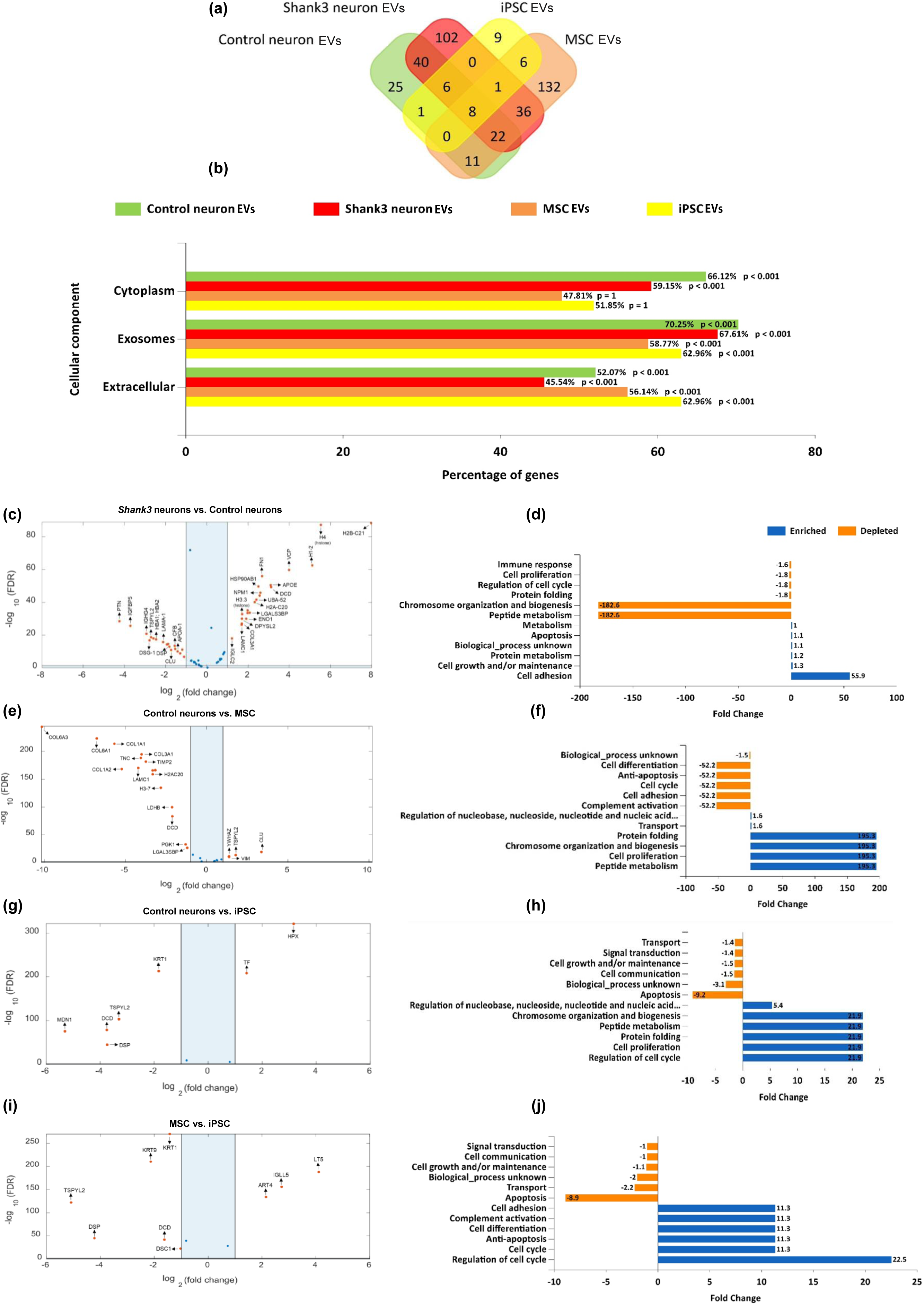
Proteomic analysis of the functions cargoes of EVs derived from control neurons, EVs derived from *Shank3* mutant neuron*s*, MSC-derived EVs, and iPSC-derived EVs. (a) A Venn diagram of the identified and overlapping proteins in EVs derived from control neurons, EVs derived from *Shank3* mutant neurons, MSC-EVs, and iPSC-EVs using LC-MS/MS. (b) The cellular component analysis of the identified proteins from four types of EVs: neuron-derived (control and *Shank3*) and stem cell-derived (MSC and iPSC) EVs. (c) A volcano plot for the differentially expressed proteins between EVs derived from *Shank3* mutant and control neurons. (d) The biological process analysis of the differentially expressed proteins between EVs derived from *Shank3* mutant and control neurons. (e) A volcano plot for differentially expressed proteins between EVs derived from control neurons and MSC-derived EVs (f) The biological process analysis of the differentially expressed proteins between EVs derived from control neurons and MSC-derived EVs. (g) A volcano plot for the differentially expressed proteins between EVs derived from control neurons and iPSC-derived EVs. (h) The biological process analysis of the differentially expressed proteins between EVs derived from control neurons and iPSC-derived EVs. (i) A volcano plot for the differentially expressed proteins between MSC-derived EVs and iPSC-derived EVs. (j) The biological process analysis of the differentially expressed proteins between MSC-derived EVs and iPSC-derived EVs. The analysis of the differentially expressed proteins was performed using FunRich proteomics software(*65*).

The *SynGO* synapse database(*32*) classification of the proteins from these EVs revealed that *Shank3* mutant neuron EVs showed significantly higher enrichment for synaptic components compared to control neuron EVs (Supplementary Fig. 4). Specifically, *Shank3* EVs were enriched for “synapse” (adjusted p = 9.92e-10), “postsynaptic ribosome” (adjusted p = 4.22e- 7), and “postsynapse” (adjusted p = 1.33e-6) while control EVs exhibited lower enrichment for these components consistent with their respective effect on the neurophysiological properties of recipient neurons. Among the stem cell-derived EVs, MSC-EVs showed higher enrichment for “synapse” (adjusted p = 9.66e-4), “synaptic cleft” (adjusted p = 1-91e-4), and “postsynaptic cytoskeleton” (adjusted p = 0.0127) compared to iPSC-EVs but less than control and *Shank3* neuron derived EVs. *Shank3* EVs carried unique protein cargoes such as *ACTB*, *Cofilin1*, and *Calsyntenin1* compared to control EVs which carried unique cytoskeletal and extracellular matrix proteins like (*ACTG1*, *TMC*, *TIMP2*, and *TPM4*) explaining their distinct neurophysiology. Interestingly, MSC-EVs had complement proteins (role in synaptic pruning) like *C1QTNF3, C1Q*, *C1R*, *C1S*, *CFD*, *CFH;* calcium activity modulators like *CACNA2D1*, *CALM3*, *CALML5*; plasticity and homeostatic regulators like *TSPYL2*, *TGFB*1, Annexins (*ANXA2*, *ANXA5*), *Fibronectin*, *Vitronectin* etc. Similarly, iPSC-EVs had synaptic regulators like *DSP*, *DSG1*, and *DSC1*, likely stabilizing synaptic junction through adhesion as well as plasticity and homeostatic regulators such as *FGF2*, *TSPYL2*, *MDK*, *GAPDH*, and *SFRP1* further explaining their neuroprotective role in *Shank3* neurons albeit with different molecular pathways.

The differential protein expression analysis between the four groups is plotted in Fig 6c,e,g,i & Supplementary Fig 5a,c as volcano plots. The biological pathways analysis of the protein cargoes for the corresponding volcano plots (Fig 6d,f,h,j and Supplementary Fig 5b,d). “Cell adhesion”, “Cell growth/ or maintenance”, “Protein metabolism”, and “Apoptosis” were among the highly enriched biological pathways in EVs derived from *Shank3* neurons compared to EVs derived from control neurons (Fig 6d). “Chromosome organization and biogenesis”, “Peptide metabolism”, “Protein folding”, “Cell proliferation” and “regulation of cell cycle” were depleted in EVs derived from *Shank3* mutant neurons compared to EVs derived from control neurons (Fig 6d). When comparing proteins extracted from MSC and iPSC-EVs compared to EVs derived from control neurons “Chromosome organization and biogenesis”, “Peptide metabolism”, “Protein folding”, and “Cell proliferation” were commonly enriched (Fig 6d,f,h). Further, when comparing MSC-EV and iPSC-EV protein cargoes, we found “Regulation of cell cycle”, “Cell cycle”, “Anti-apoptosis”, “Complement activation”, and “Cell adhesion” to be enriched while “Apoptosis”, “Transport”, “Cell growth/ or maintenance”, “Signal transduction” were down-regulated (Fig 6i,j).

### *Shank3*-knockout (KO) mice do not exhibit emotional state preference which is restored by intranasal iPSC-EV treatment

Since human individuals diagnosed with ASD are known to exhibit impaired emotion recognition(*33–35*), we examined whether *Shank3*-KO mice, a well-established genetic mouse model for ASD(*36*), are also impaired in their emotion recognition ability. To that end, we employed the emotional state preference (ESP) task (Fig 7g), previously reported by us and others(*37*, *38*), in which subject mice are exposed simultaneously to a stressed and a naïve conspecific. We started by comparing *Shank3*-KO mice behavior in the ESPs task, to their behavior in the social preference (SP; social vs. object stimuli, Fig 7a) and sex preference (SxP; male vs. female stimuli, Fig 7d) tasks(*39*), which do not rely on the emotional state of the stimulus animal. We found that both wild-type (WT) and *Shank3*-/- (KO) male mice exhibited a similarly clear preference in the SP and SxP tasks (Fig 7b, e). No difference between the two genotypes was found in the distance traveled during any of the tasks (Fig 7c, f, i), indicative of no change in their motor activity. In contrast, only WT mice, but not *Shank3*-KO mice, exhibited a significant preference in the ESPs task (Fig 7h; Supplementary Video 1,2) (WT: stressed stimulus: 101.4±8.1, Naïve stimulus: 75.1±5.2; KO: stressed stimulus: 114.3±6.9, Naïve stimulus: 96.6±5.9); Significant main effects were revealed by a two-way Mixed-Model (MM) ANOVA for stimulus (*p<*0.01) and genotype (*p<*0.05). Thus, *Shank3*-KO mice are impaired specifically in their ESP behavior.

**Figure 7.**
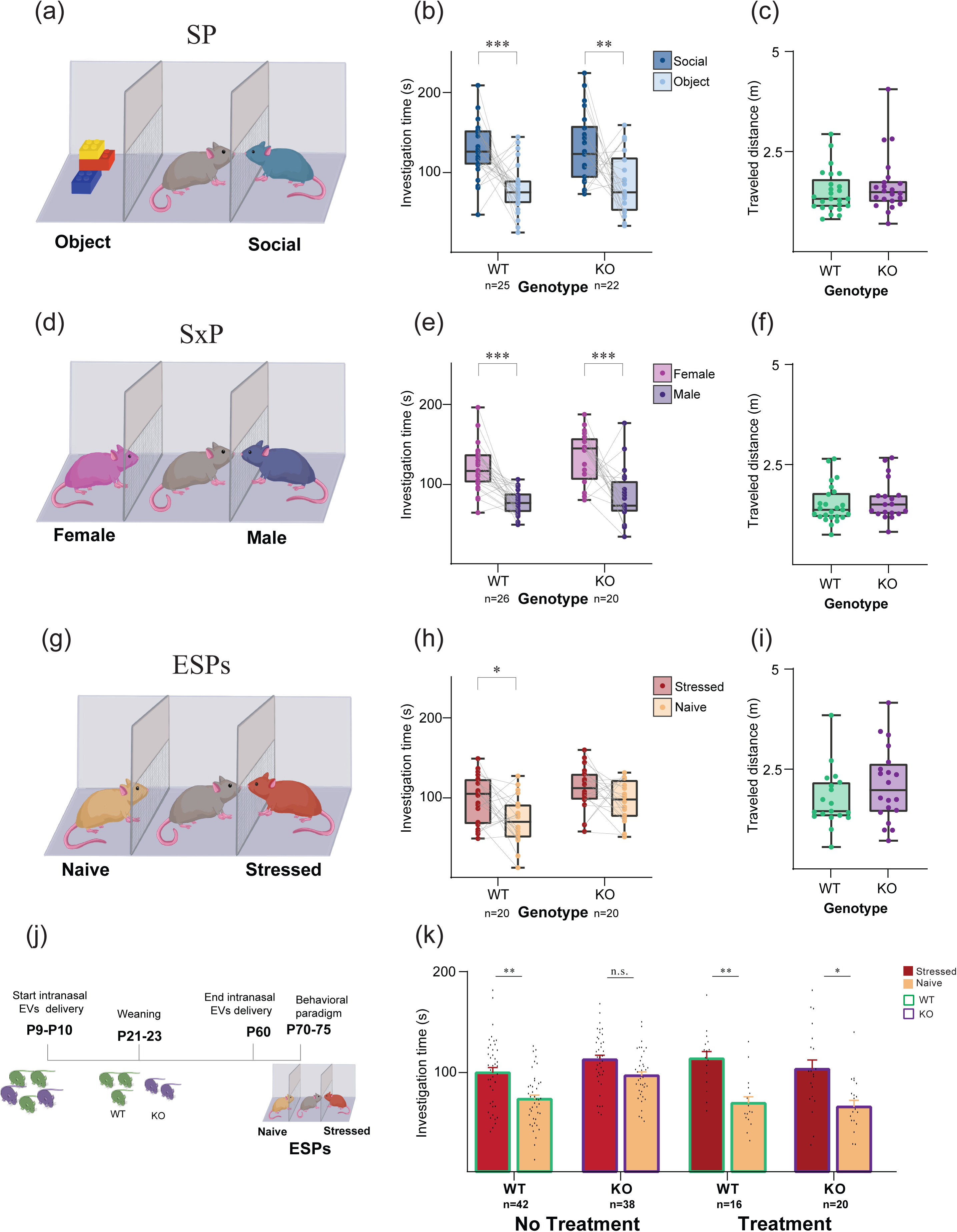
Intranasal treatment with iPSC derived-EV restores the behavioral deficit exhibited by *Shank3-KO* mice in emotion recognition. (a) Schematic representation of the Social Preference (SP) task. (b) Mean (±SEM) time dedicated by wild-type (WT; left) and *Shank3*-KO (KO; right) male mice for investigating the social (dark blue) or object (light blue) stimulus during the SP task (sample size denoted below bars). A significant main effect was found in two-way mixed-model (MM) ANOVA for stimulus (*p<*0.001). (c) Mean (±SEM) distance traveled by WT (left) and KO (right) subjects during the SP task. (d-f) As in a-c, for the sex preference (SxP) task. Significant main effects were revealed by a two-way MM ANOVA for stimulus (*p<*0.001) and genotype (*p<*0.05). (g-i) As in a-c, for the emotional state preference (ESPs) task. Significant main effects were revealed by a two-way MM ANOVA for stimulus (*p<*0.01) and genotype (*p<*0.01). (j) A schematic description of the experiment, with the ESP task in adulthood following intranasal EV treatment from early postnatal to juvenility of WT and *Shank3*-KO littermates. (k) Mean (±SEM) time dedicated by WT and KO mice (sample size denoted below bars), without (left) or with (right) intranasal EV treatment, for investigating the naive (orange) or stressed (red) stimulus during the ESPs task. Significant main effects were revealed by a three-way MM ANOVA for stimulus (*p<*0.001), interaction between genotype and treatment (*p<0.01*), and interaction between stimulus and treatment (*p<*0.05). *\*p<0.05, **p<0.01, ***p<0.001*, post-hoc paired t-test with Bonferroni correction for multiple comparisons following the identification of main effects by ANOVA.

To examine if treating *Shank3*-KO animals with intranasal administration of EVs affects the ESP behavior, we applied the treatment between P9-10 days to P60 days (see methods) and examined the behavior of the animals on P70-75 days (Fig 7j; Supplementary Video 3). We found that, unlike untreated *Shank3*-KO mice, which exhibited impaired ESP behavior, EV-treated animals showed normal ESP behavior (Fig 7k, Supplementary Video 4). Accordingly, a three-way ANOVA analysis of the effects of Stimulus, Treatment, and Genotype yielded a significant interaction between Treatment and Genotype (*p<*0.01), with a significant difference between the genotypes in the condition of no treatment (*p<*0.05; post-hoc paired t-test with Bonferroni correction for multiple comparisons) and no significant difference between them after treatment. Thus, the intranasal application of EVs during the early postnatal to juvenility managed to restore the normal ESP behavior in *Shank3*-KO mice in adulthood.

## DISCUSSION

The absence of distinct pathological hallmarks in ASD has been a limiting factor in detecting possible effects of therapeutic drugs and molecules, due to the lack of suitable read-out modalities. ASD, unlike several neurodegenerative diseases, such as AD and PD, does not exhibit protein aggregation pathologies like Amyloid-beta plaques or *Lewy bodies* in neurons that can serve as biomarkers and targets for novel therapeutics. Our current study along with previous studies has identified the accelerated maturation in iPSC-derived neurons of individuals with ASD with *Shank3* and other ASD-associated mutations as one potential endophenotype and read-out(*15*, *40–42*). This accelerated maturation was measured by increased Na+ currents, increased excitability, and increased EPSC rate early in the differentiation(*15*, *43*), as well as in transcriptomic and morphological analysis(*40*). Further, evidence for increased neuronal connections at the early stages of development was observed in an fMRI study of children with ASD that found brain hyperconnectivity correlated with the severity of social dysfunction(*44*). Additionally, increased spine density was found in the post-mortem brain samples of children with ASD at the age of 2-9 years compared to age-matched controls(*10*, *45*). These findings point to increased synaptic connections in ASD and thereby correlate to the increased EPSC frequency that we have observed in patients’ iPSC-derived neurons with a *Shank3* mutation.

EVs hold great potential in translational research especially in therapeutics and diagnostics due to their role in inter-cellular communication(*19*). We started the study by switching the EVs isolated from healthy and *Shank3* mutant neurons. Surprisingly, treating control neurons with EVs derived from *Shank3* mutant neurons was sufficient to induce ASD phenotypes in control neurons. However, the EVs derived from control neurons did not rescue the phenotypes of *Shank3* mutant neurons. Recent studies have demonstrated that the source or parent cells from which the EVs are produced greatly influence the cargo they carry to the recipient cells(*18*, *46*, *47*). Our results have thus important implications for understanding the pathophysiology of ASD as well as cellular communications mediated via EVs in the human central nervous system. We demonstrate the effect of EVs isolated from different cells on neuronal function and connectivity. Interestingly, in this study, we show for the first time that EVs derived from MSCs and iPSCs rescue abnormal neurophysiological phenotypes of *Shank3* mutant human neurons, due to the presence of plasticity and homeostatic regulator cargoes. We additionally demonstrate that *Shank3* phenotypes can be transferred to control neurons via EVs derived from the mutant neurons driven by enriched synaptic protein cargoes compared to other EVs. The trans-synaptic mode of communication via EVs has been demonstrated previously for proteins and cell-signaling factors in the neuromuscular junction in *Drosophila* (*48*, *49*). Hence, the transfer of intrinsic properties of excitability of ASD neurons through our results further reinforces an alternative mode of communication between human neuronal cells via extracellular vesicles.

Despite the clear genetic basis of *Shank3*-related pathologies, there are currently no approved drugs specifically targeting the broader spectrum of core ASD symptoms in patients with *Shank3* mutations. Given the ability of stem-cell-derived EVs to cross the blood-brain barrier and being minimally invasive(*50*), they offer hope as a possible therapeutic approach for neurological disorders(*27*). We tested the effects of purified EVs from MSCs and iPSCs on *Shank3* cortical neurons to better understand their impact on neuronal intrinsic properties and synaptic connections. Some beneficial properties of MSCs have been attributed to their paracrine effect mediated via EVs(*51*). In our study, we found that MSC-derived EVs rescue the accelerated maturation of *Shank3* mutant neurons. Notably, MSC-derived exosomes were previously shown to rescue social interaction and communication deficits in mouse models of autism(*30*, *52*). Although MSC-derived EVs have shown promise for clinical applications, certain limitations in scaling up MSC cultures for EV production present significant challenges(*53*). MSCs, for example, have a limited proliferative capacity and have been found to exhibit chromosome variability, cellular senescence, and molecular changes after 4-5 passages in *in vitro* cultures leading to higher costs for EV production(*54*, *55*). In addition, to extract MSCs, a donor needs to undergo an invasive procedure. Furthermore, the donor source and isolation methods can affect the quality and potency of MSCs and thus the EVs produced from them(*56*). Due to their pluripotent nature, iPSCs offer solutions to these challenges with their almost unlimited proliferative capacity, comparatively lesser chromosomal variability, and higher consistency in *in vitro* conditions facilitating large-scale production for therapeutic use(*57*, *58*). iPSCs can be produced with minimal stress to the donor (a blood or even a urine sample).

Similar to MSC-derived EVs, we found that iPSC-derived EVs rescued the accelerated maturation in *Shank3* mutant neurons in *in vitro* cultures. Additionally, our results are further strengthened by the restoration of emotional state recognition abilities in an *in vivo* mice model of ASD after iPSC-derived EV treatment. We prioritized the timeframe of treatment for *Shank3B*-/- mice from the early postnatal period which coincides with the critical period of brain plasticity enabling substantial improvement in behavioral deficits. Furthermore, we employed the intranasal method of EV delivery which is non-invasive; has minimal systemic exposure, and is easier for clinical applications, especially for pediatric populations.

Proteomic analysis of EVs revealed distinct protein cargo profiles, especially differences were observed between EVs derived from stem cells (MSCs and iPSCs) and EVs derived from neurons (control and *Shank3* mutant). Importantly, although MSC-derived EVs and iPSC-derived EVs had similar protective effects on the *Shank3* mutant neurons, the protein cargo was not identical with more than half of the unique protein content and higher enrichment of synaptic pruning (complement proteins) and calcium signalling modulators in MSC-EVs; Plasticity regulators and extracellular matrix proteins were more abundant in iPSC-EVs. The MSCs are multipotent and mesodermal while iPSCs can differentiate into the three germ layers. Hence, the proteomic profile of the EVs likely reflects the underlying physiological properties of the parent cells(*46*). Interestingly, we also found that MSC-derived EVs carry a larger protein cargo load compared to iPSC–derived EVs. However, towards the development of EV-based therapies, we believe it would be more practical to engineer EVs with minimal cargo while maintaining comparable efficacy to those with a larger cargo load given the complexity of intermolecular interactions.

In conclusion, using accelerated maturation as an endophenotype for ASD-associated mutations, we provide evidence that EVs can affect the intrinsic and network properties of *Shank3* mutant neurons as well as offer a potential therapeutic approach for modulating behavioral deficits in neurodevelopmental disorders. The ability of EVs to transfer pathological phenotypes between neurons reinforces their significance in intercellular communication and highlights the need for further research into their therapeutic applications. Future studies should focus on characterizing the specific mechanism by which functional cargoes are loaded in EVs from different cell types. This will be essential for developing targeted EV-based therapies for ASD and related neuropsychiatric disorders.

## METHODS

### Ethics approval

All iPSC experiments were performed following the relevant guidelines set by the institutional review board (IRB), University of Haifa, Israel, and approved by the IRB. The animal experiments were performed according to the National Institutes of Health guide for the care and use of laboratory animals and approved by the Institutional Animal Care and Use Committee of the University of Haifa (Ethic number: UoH-IL2203-141-4, 1076U).

### iPSC culture and generation of cortical Neural Progenitor cells (NPCs)

iPSCs carrying a (c.3679insG) *Shank3* mutation and the control iPSCs were previously(*14*) generated from a female child and its unaffected mother’s fibroblasts respectively. iPSCs were cultured and maintained using mTesR PLUS media. Cortical NPCs were generated using a previously described protocol (*15*, *59*, *60*). In brief, iPSCs were grown to ∼80% confluency, then dissociated using Dispase (StemCell Technologies, Cat #07923) and plated onto low-adherence plates (Falcon, 351007) in mTeSR medium with ROCK inhibitor, allowing for the formation of embryonic bodies (EBs). The media was changed on the next day to mTeSR PLUS without ROCK inhibitor. In the following 10 days, the cells were fed with EB media containing: DMEM/F12 with Glutamax (1:100, Gibco, Cat #35050038), B27 with Retinoic Acid (1:50, Gibco, Cat #17504044), N2 supplement (1:100, Gibco, Cat #17502048), and 0.1uM LDN- 193189 Hydrochloride (Biogems, Cat #1066208). The EBs were plated onto poly-L-ornithine/laminin (Sigma-Aldrich, Cat # P3655, R&D Systems, Cat #3400-010-03) coated six-well dishes in DMEM/F12 plus N2, B27, and laminin for the following 7 days to allow the formation of neural rosettes. The rosettes were selected based on their morphology and were manually picked, dissociated (with Accutase (StemCell Technologies, Cat #07920)) and plated onto poly-l-ornithine/laminin-coated plates in neural progenitor cell (NPC) medium containing: DMEM/F12 with Glutamax (1:100), B27 supplement with RA 50X (1:50), N2 supplement 100X (1:100), laminin (1 mg/ml), and 20 ng/ml bFGF. Full media change was performed every second day for the following 7-10 days until full confluency. Cortical NPCs thus generated were passaged and expanded.

### Differentiation of cortical NPCs into cortical neurons

Cortical NPCs after 3 to 4 passages were differentiated into cortical neurons by a differentiation medium containing: DMEM/F12, N2, B27 with retinoic acid, Glutamax, 200 picoMol L-Ascorbic Acid (Biogems, Cat #5088177), 500 µg/ml Dibutyryl-cAMP (Adooq, Cat #A15914- 5), 1 mg/ml laminin, 20 ng/ ml Brain-derived neurotrophic factor (BDNF) (Peprotech, Cat #AF-450-02) (20 ng/ ml), and 20 ng/ml Glial cell line-derived neurotrophic factor (GDNF) (Peprotech, Cat # 450-10) for 10 days. Between days 11 and 14, the cells were dissociated again, plated on 24 or 48 well coverslips and then fed with Brainphys medium with B27 with retinoic acid, N2, L-ascorbic acid (200 picoMol), cyclic AMP (500 µg/ml), BDNF (20 ng/ml), GDNF (20 ng/ml), and laminin (1 mg/ml). The neurons were differentiated for up to 5 weeks and the experiments were performed starting 4 weeks of differentiation.

### Extracellular vesicle Isolation and treatment *in vitro*

#### EVs from Control and Shank3 neurons

The EVs were purified from iPSC-derived Control and *Shank3* mutant cortical neurons using differential centrifugation followed by an ultra-centrifugation method as described previously (*61*). Briefly, the iPSC-derived cortical neurons were cultured and maintained in cortical neuronal differentiation media, as described above, devoid of any exogenous source of extracellular vesicles. The culture media collected from the iPSC-derived cortical neurons were centrifuged for 10 min. at 300*g* at 4 °C. The pellet was discarded and the supernatant was recovered and re-centrifuged for 10 min. at 2000*g at* 4 °C. The pellet of dead cells was then discarded and the supernatant was collected again and centrifuged for 30 min. at 10,000*g* (4 °C) to remove the cell debris. The supernatant (free of dead cells and cell debris) was filtered through a 0.22-μm filter and ultra-centrifuged for 70 min at 100,000*g* (4 °C). The pellet containing the EVs and proteins was washed in 0.22-μm filtered PBS and then ultra-centrifuged again for 70 min. at 100,000*g* (*4 °C*). Finally, the pellet containing the purified EVs was re- suspended in 100-200 μl of filtered and sterilized PBS.

#### EVs from MSCs and iPSCs

Human bone marrow MSCs (Lonza, Cat #PT-2501) were cultured in DMEM/F12 medium, supplemented with 1% 200mM L-glutamine (Thermo Scientific, Cat # 25030081), 1% MEM-Eagle nonessential amino acid (Thermo scientific Cat # 11140050), 0.04% heparin (Sigma, Cat # H3393), and 10% extracellular vesicle-free platelets lysate (Rabin Medical Center, Israel) for 3 days. Conditioned media was collected from the cells and extracellular vesicle isolation was done using a differential centrifugation method followed by ultracentrifugation as described previously(*52*, *61*). For iPSC-EV, iPSCs from a healthy individual were cultured as described above and the EVs were isolated by the same process described above after the collection of the conditioned media from the culture of control healthy iPSCs.

#### Synthetic liposome production

Synthetic liposomes were produced as described previously(*62*) using a Nanoassemblr (Precision nanosystem). Synthetic liposome treatment was performed two times at a similar time point as the extracellular vesicle switching experiment.

#### Extracellular vesicle treatment in vitro

EVs from control and *Shank3* cortical neurons were switched two times –one during the first 7 days of differentiation and the second approximately 14-15 days of differentiation. EVs from MSCs and iPSCs and synthetic liposomes were applied three times on *Shank3* cortical neurons. First within 3-4 days of differentiation, second after 14-15 days of differentiation, and third after 23-24 days of differentiation. 2 µl of 2-3*10^10^ particles/ µl were added in each well containing 200-300K cells. The electrophysiology was performed within a week of the last treatment of EVs at approximately 29-30 days of the differentiation.

### Extracellular vesicle characterization, labeling, and uptake

EVs were characterized using Zetasizer (Malvern Panalytical) for charge, particle size, and concentration using dynamic light scattering (DLS) and electrophoretic light scattering methods (ELS). Briefly, for size and concentration measurements, 10 ul of EVs were diluted in 990 ul of 0.22 µm filtered PBS (1X) in disposable size cuvettes (DTS0012) and measurements were done using the Ultra-pro ZS Xplorer software. For charge, 10 ul of EVs were diluted in 90 ul of PBS (1X) and 900 ul of MilliQ water in a glass ZP cuvette (DTS1070). The measurements were done similarly in size and concentration between all the samples. Further, the EVs were characterized using proteomics for the protein markers as described below. EVs were labeled using PKH-67 (Merck, MIDI67) dyes as described previously(*30*, *52*). The labeled EVs were imaged within 24 hours of incubation and were observed to be up-taken by the neurons.

### Electrophysiology

Whole-cell patch-clamp recordings were performed in *Shank3* mutant cortical neurons, and from control neurons that were treated with EVs isolated from neurons, MSCs, and iPSCs based on previously described methods(*63*, *64*) 4-5 weeks after the start of the differentiation. Culture coverslips were placed inside a recording chamber filled with HEPES-based artificial cerebrospinal fluid (ACSF) containing (in mM): 10 HEPES)Sigma-Aldrich, Cat #H0887- 100ML(, 139 NaCl (Sigma-Aldrich, Cat #S9888), 4 KCl (Sigma-Aldrich, Cat #P3911), 2 CaCl2 (Sigma-Aldrich, Cat #223506), 10 D-glucose (Sigma-Aldrich, Cat # G7021), and 1MgCl2 (Merck, Cat #7791-18-6) (pH 7.5, osmolarity was adjusted to 310 mm). Fire-polished borosilicate glass capillaries (Sutter Instrument, Cat #BF150-75-10) were pulled (tip resistance of about 9–12 MΩ) using a (P1000, Sutter Instrument, Novato, CA, United States) and filled with an internal solution containing (in mM): 130 K-gluconate, 6 KCl, 4 NaCl, 10 Na-HEPES, 0.2 K-EGTA, 0.3 GTP, 2 Mg-ATP, 0.2 cAMP, 10 D-glucose, 0.15% biocytin and 0.06% rhodamine (pH 7.5, osmolarity adjusted to 290–300 mOsm). All measurements were done at room temperature using a patch clamp amplifier (MultiClamp 700B, Molecular Devices, San Jose, CA, United States), connected to a digitizer (Axon Digidata 1550B, Molecular Devices, San Jose, CA, United States), and controlled by MultiClamp 700B Commander and pCLAMP 11 software. Data were acquired at a sampling rate of 20 kHz and analyzed using Clampfit-10 and the software package MATLAB (release 2014b; The MathWorks, Natick, MA, USA).

### Analysis of electrophysiological recordings

The acquired data was analyzed based on previously described methods (*63*) using custom-written MATLAB scripts. Briefly: *Sodium, fast, and slow potassium currents*. Neurons were held in voltage clamp mode at −60 mV, and voltage steps of 400 ms were performed in the −100 to 90 mV range The sodium (Na+) current was computed by subtracting the sodium current after stabilization from the lowest value of the inward sodium current. The fast potassium (K+) currents were measured by the maximum outward currents that appeared within a few milliseconds after a depolarization step. The slow K+ currents were measured at the end of the 400 ms depolarization step. A one-way ANOVA test was performed for the statistical analysis between the groups. Currents were normalized by the cells’ capacitance. The capacitance was measured following the instructions in Clampex SW.

*Evoked action potentials* (APs). Neurons were held in current clamp mode at −60 mV with a constant holding current. Following this, current injections were given in 3 pA steps with 400 ms duration, starting 12 pA below the steady-hold current. A total of 38 depolarization steps were given. The total evoked action potential was the total number of action potentials that were counted in the 38 depolarization steps. Non-parametric statistical tests (Mann-Whitney U test) were performed for comparisons between the groups since the data were not normally distributed.

*Synaptic currents analysis*. The excitatory postsynaptic currents (EPSCs) recordings were done by holding the neurons at −60 mV in voltage-clamp mode. The mean and standard error (SE) of EPSC amplitudes for each active cell were calculated. The cumulative distribution of EPSC amplitude was calculated for each group and condition. The frequency of the events for each cell was calculated by dividing the number of events by the duration of the recording (non-active cells were included with an event rate of 0 Hz). The mean rates and standard errors of EPSC frequencies of all the cells in each group were computed. Non-parametric statistical tests (Mann-Whitney U test) were performed for comparisons.

### Proteomics

For Proteomics, the LC-MS/MS facility at the *Smoler Proteomics Centre* facility at Technion, Haifa, Israel was availed. Briefly, the EVs ∼50 ul (10^14^- 10^15^ particles/ul) were lysed in RIPA buffer and the extracellular vesicle-RIPA buffer mix was then incubated on ice for 20-30 mins for the extraction of proteins. The sample was then centrifuged at 16000g for 10 min. at 4 °C. The cleared lysate (supernatant) was used for protein estimation using the Pierce ^TM^ (BCA) assay kit (Thermo Scientific, 23225). The extracted protein was trypsinized and the samples were subjected to the LC-MS/MS analysis using the Q Exactive HF mass spectrometer. The data was analyzed using the Proteome Discoverer 2.4 software. The graphs for proteomics data analysis were made using the Funrich analysis tool (*65*, *66*). The volcano plots of differentially expressed proteins between the various EVs were plotted using MATLAB (*67*).

### Immunocytochemistry (ICC)

Cells on coverslips were fixed in 4% paraformaldehyde for 15 min. and then washed 3 times with DPBS for 5 min. In each step, they were blocked and permeabilized in PBS containing 0.1–0.2% Triton X-100 and 10% horse serum. Next, the coverslips were incubated with primary antibodies; for Neurons: chicken anti-MAP2 (Abcam, ab92434, 1:500), rabbit anti-TBR1 (Abcam, ab183032, 1:300), anti-CTIP2 (Abcam, ab18465, 1:500) in the blocking solution overnight at 4 °C. On the next day, the coverslips were washed in DPBS and incubated with DAPI (Abcam, ab228549, 1:2500) and the corresponding secondary antibodies for 60 min. at room temperature. Then, the coverslips were washed three times, mounted on glass slides using Fluromount-G (mounting medium), and dried overnight while being protected from the light. Microscopy was performed using a Leica THUNDER imager and ZEISS LSM 980 with Airyscan 2. The images were analyzed using ImageJ and Imaris software.

### Animals

Social stimuli: Social stimuli were ICR strain adult male or female mice (8-18 week-old) in the Emotional Stress Preference (ESP) and Sex preference (SxP) tasks, or juvenile ICR male mice (3-6 week-old) in the Social Preference (SP) task.

*Shank3* subjects: B6.129-*Shank3^tm2Gfng^*/J mutant mice (*Shank3B*)(*36*) were crossed by us for five generations with ICR mice to create *Shank3B* mutant mice with a B6; ICR background. All *Shank3B*^−/−^ (KO) and wild-type (WT) mice used in this study were obtained by crossing male and female *Shank3B*^+/−^ mice born with the expected Mendelian frequencies. *Shank3B* subjects with KO or WT genotypes were used for behavioral testing at 8-18 weeks of age.

Genotyping: Ear tissue samples were collected from offspring mice at 21 days of age for genotyping by polymerase chain reaction (PCR) using the following primers: WT forward primer: GAGCTCTACTCCCTTAGGACTT; WT reverse primer: TCCCCCTTTCACTGGACACCC, yielding a 250 bp band; Mutant forward primer: GAGCTCTACTCCCTTAGGACTT; Mutant reverse primer: TCAGGGTTATTGTCTCATGAGC, yielding a 330 bp band. Maintenance: All animals were kept in groups of 2-5 sex-matched mice per cage at the animal facility of the University of Haifa under veterinary supervision, in a 12-hour light/12-hour dark cycle (lights on at 9 PM), with *ad libitum* access to standard chow diet (Envigo RMS, Israel) and water.

### Intranasal extracellular vesicle treatment to the WT and KO mice

The intranasal extracellular vesicle treatment was started on postnatal days 9-10 and was administered every other day till the postnatal day 60 (2 months). The mouse pups were given 2 µl of 2-3*10^10^ particles/µl/mouse till postnatal 21-23 days (until weaning). After weaning, 5 µl of 2-3*10^10^ particles/µl was administered per mouse intranasally (Supplementary Video 3).

### Experimental setups

Behavioral experiments were conducted in the dark phase of the dark/light cycle in a sound attenuated chamber, under dim red light. The experimental setup(*68*) consisted of a black Plexiglas arena (37 X 22 X 35 cm) placed in the middle of an acoustic cabinet (60 X 65 X 80 cm). Two Plexiglas triangular chambers (12 cm isosceles, 35 cm height), into which the stimulus (a plastic toy in the case of object stimulus, or an animal in all other cases) could be introduced, were placed in two randomly selected opposite corners of the arena. A metal mesh (12 X 6 cm, 1 X 1 cm holes) located at the bottom of the triangular chamber allowed direct interaction with the stimulus through the mesh. A high-quality monochromatic camera (Flea3 USB3, Flir) equipped with a wide-angle lens was placed at the top of the acoustic chamber and connected to a computer, enabling a clear view and recording (∼30 frames/s) of subject behavior using commercial software (Fly Capture2, FLIR).

### Behavioral paradigms

*Social preference (SP):* Subjects were taken from their home cage and placed in an arena with empty chambers for a 15-minute habituation period. Throughout this time, social stimuli were placed in their chambers near the acoustic cabinet for acclimation. After habituation, the chamber containing the social and object stimuli was diagonally placed at opposite ends of the arena randomly, and the SP task was conducted for 5 min.

*Sex Preference (SxP):* The SxP task consisted of 15 minutes of habituation to the arena with empty chambers, followed by exposing the subject to both adult male and female social stimuli located in individual chambers found at opposite corners of the arena for 5 minutes.

*ESP task*: The task consists of 15 min of habituation followed by a 5-minute period in which the subject mouse was introduced to a naive stimulus animal and to another stimulus animal that had been constrained for 15 minutes in a 50 ml tube pierced with multiple holes for ventilation. Each “stressed” stimulus animal was used as a stimulus for only two consecutive sessions (Supplementary Videos 1,2, and 4).

### Quantification & statistical analysis of behavioral analysis

#### Tracking software and behavioral analyses

All recorded video clips were analyzed using TrackRodent (https://github.com/shainetser/TrackRodent)(68). Behavioral analysis was conducted as previously described(*68*).

#### Statistical analysis

The sample size for all behavioral experiments was based on previously published power calculations(*68*). All statistical tests were performed using MATLAB 2024a. The Shapiro-Wilk test was used for verifying the normal distribution of the dependent variables and Levene’s test for homogeneity of variances. A two-tailed paired t-test was used to compare different conditions or stimuli for the same group, and a two-tailed independent t-test was used to compare a single variable between distinct groups. For comparison between multiple groups and parameters, a mixed model (MM) analysis of variance (ANOVA- either two-way or three-way) was applied to the data. All ANOVA tests were followed, if main effects or interactions were found, by a *post-hoc* Student’s t-test with Bonferroni correction.

## ACKNOWLEDGMENTS

The Zuckerman STEM leadership program and Israel Science Foundation grants - 1994/21 and 3252 /21 for Prof. Shani Stern. The Maof Fellowship for the Integration of Outstanding Faculty, Council for Higher Education (2023-2025) for Ahmad Abu-Akel. IBBRC - The Integrated Brain And Behavior Research Centre and Graduate Studies Authority, Bloom Graduate School, University of Haifa, Israel, for supporting our research. The schematic images in the manuscript were created with Biorender.com

## AUTHOR CONTRIBUTIONS

**AC** and **IR** (co-first authors) contributed equally to the work**. AC**- Experiments-Cell culture, EV isolation, labeling, quantification, EV treatment *in vitro* and *in vivo*, Immunocytochemistry, Proteomics Data Analysis, Methodology design, Drafting the original manuscript. **IR-** Electrophysiology, iPSC culture, EV quantification & *in vivo* treatment, Methodology design, help in drafting the original manuscript. **YH**- Electrophysiology Data analysis. **SN**: Behavioral experiments design and analysis; Drafting manuscript **AS**- Proteomics Data analysis. **TS:** Behavioral experiments **WAR**: cell culture experiments. **LS & BS**: Help in imaging; Resource availability for imaging **AAA**: Resource availability and editing manuscript **DO**-Resource availability for EVs. **AZ**-Resource availability for EV quantification. **SW** (corresponding author): Supervision, Conceptualisation & Methodology design-*in vivo* experiments, Editing-original draft. **SS** (corresponding author)- Supervision, Conceptualization, Methodology design, MATLAB codes for Data Analysis, Editing –original manuscript. All authors reviewed the manuscript.

## DECLARATION OF INTERESTS

### Competing Interests

The Authors declare no competing financial or non-financial Interests.

## DATA AND CODE AVAILABILITY

The data that support the findings of this study are available upon reasonable request from the authors. The MATLAB codes used for mouse behavioral data analysis are provided in the GitHub link in the methods. The codes for electrophysiology data analysis and figure generation are available upon reasonable request from the authors.

## Supplementary Materials

**Supplementary Table 1-** Protein Markers according to *MISEV*(*29*) guidelines for all four type of EVs isolated

**Supplementary Figure 1-** (a) Immunocytochemistry images of the individual channels of *TBR1* (green), *CTIP2* (red), *MAP2* (orange), and DAPI (blue) for (i) untreated control neurons, (ii) control neurons treated with EVs derived from *Shank3* mutant neurons, and (iii) control neurons treated with synthetic liposomes. The EVs were added at two-time points (see methods) (The merged images are shown in Fig 2b); Scale bar, 20 µM. (b) The quantification of the *TBR1+* and *CTIP2+* neurons among *MAP2+* neurons in (i) untreated control neurons, and (ii) control neurons treated with EVs derived from *Shank3* neurons.

**Supplementary Figure 2-** (a) Immunocytochemistry images of *TBR1* (green), *CTIP2* (red), *MAP2* (orange), and DAPI (blue) for untreated *Shank3* mutant neurons and *Shank3* mutant neurons treated with EVs derived from control cortical neurons. The EVs were added at two-time points (see methods) (The merged images are represented in Fig 3a); Scale bar, 20 µM. (b) Immunocytochemistry images of the individual channels of *TBR1* (green), *CTIP2* (red), *MAP2* (orange), and DAPI (blue) for untreated *Shank3* mutant neuron*s* and *Shank3* mutant neurons treated with MSC-derived EVs. The EVs were added three times during the differentiation (see methods) (The merged images are shown in Fig 4c); Scale bar, 20 µM.

**Supplementary Figure 3 -** (a) Immunocytochemistry images of the individual channels of *TBR1* (green), *CTIP2* (red), *MAP2* (orange), and DAPI (blue) for untreated *Shank3* mutant neurons and *Shank3* mutant neurons treated with iPSC-derived EVs. The EVs were added three times during the differentiation (see methods) (The merged images are represented in Fig 5c); Scale bar, 20 µM. (b) The quantification of the *TBR1+* and *CTIP2+* neurons among *MAP2+* neurons in (i) *Shank3* mutant neurons, (ii) *Shank3* mutant neurons treated with EVs derived from control cortical neurons, (iii) *Shank3* mutant neurons treated with MSC-derived EVs, and (iv) *Shank3* mutant neurons treated with iPSC-derived EVs.

**Supplementary Figure 4 -** (a) *SynGO* cellular and biological component pathways for EVs derived from control neurons showing enrichment for synaptic proteins (b) Similarly, for EVs derived from *Shank3* mutant neurons showing higher enrichment for synaptic proteins. (c) Similar plots for MSC-EVs show enrichment for synaptic protein but lesser than neuron-derived EVs. (d) The iPSC-EVs SynGO plots do not show high enrichment for synaptic proteins exclusively.

**Supplementary Figure 5 -** (a) A volcano plot displaying the differentially expressed proteins identified between EVs derived from *Shank3* mutant neurons and MSC-derived EVs. (b) The biological process analysis of the differentially expressed proteins between EVs derived from *Shank3* mutant neurons and MSC-derived EVs. (c) A volcano plot displays the differentially expressed proteins identified between EVs derived from *Shank3* mutant neurons and iPSC-derived EVs. (d) The biological process analysis of the differentially expressed proteins between EVs derived from *Shank3* mutant neurons and iPSC-derived EVs.

**Supplementary Video 1**- A representative ESP test video of the WT ICR mouse. The mouse was presented to neutral and stressed mice (stimuli) for 5 minutes after a 15-minute habituation period (for details, see methods). The video has been speeded up 5X times.

**Supplementary Video 2**- A representative ESP test video of the *Shank3* KO ICR mouse. Other details same as in Supplementary Video 1.

**Supplementary Video 3**- A video showing intranasal administration of iPSC EVs to *Shank3* ICR mice. The intranasal treatment was started at P9-P10 days-old mice pups and was continued till P60, every other day. The ESP behavioral test was done after P60 (also see methods).

**Supplementary Video 4**- A representative ESP test video of the *Shank3* KO ICR mouse after intranasal iPSC EV treatment. Other details are the same as in Supplementary Videos 1 and 2.

## Notes

### Competing Interest Statement

The authors have declared no competing interest.

### Summary of Updates

Fig 7 added; supplementary files updated Classification of EVs according to MISEV guidelines Change in Title and abstract and new list of authors who have contributed

