## Supplementary Figures 1-5 for "Extracellular Vesicles from Stem cells Rescue Cellular Phenotypes and Behavioral Deficits in SHANK3-Associated ASD Neuronal and Mouse Models"

#### Supplementary Table 1

| EV origin | Protein markers |
|---|---|
| Control neuron EVs | ANXA2, ACTG1, LGALS3BP, COL*, TUBB*, GAPDH, FN1 |
| *Shank3* neuron EVs | B2M , CD81, PDCD6IP, LGALS3BP, ACTB, COL*, TUBB*, TGFB1,FN1 |
| MSC EVs | B2M, CD63, CD81, CHMP4A, NT5E, PDCD6IP,COL*, TUBB*,FN1 |
| iPSC EVs | ANXA1, DSG1, DSC1, GAPDH, FGF2 |

#### Supplementary Figure 1

(a)

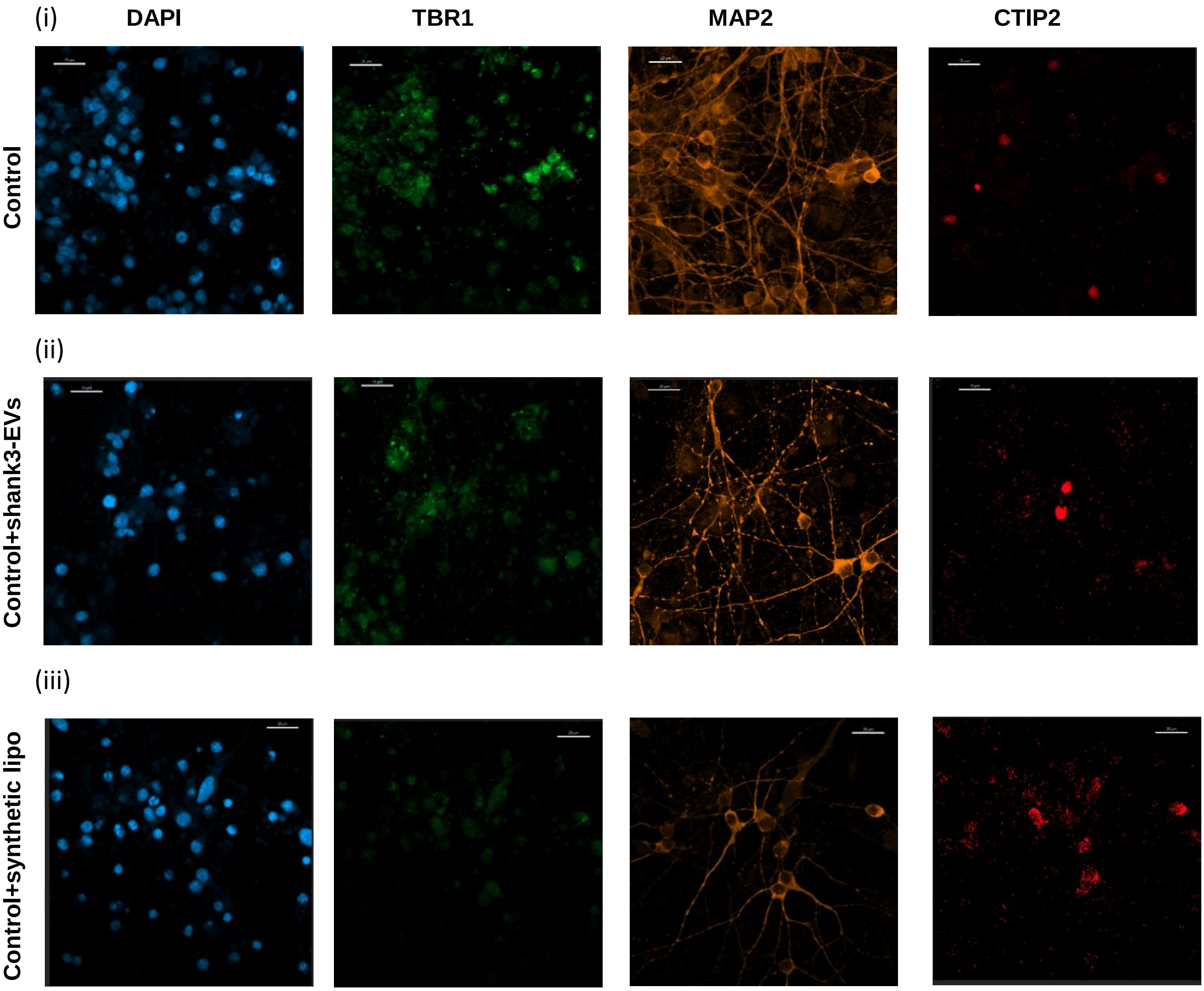


(b)

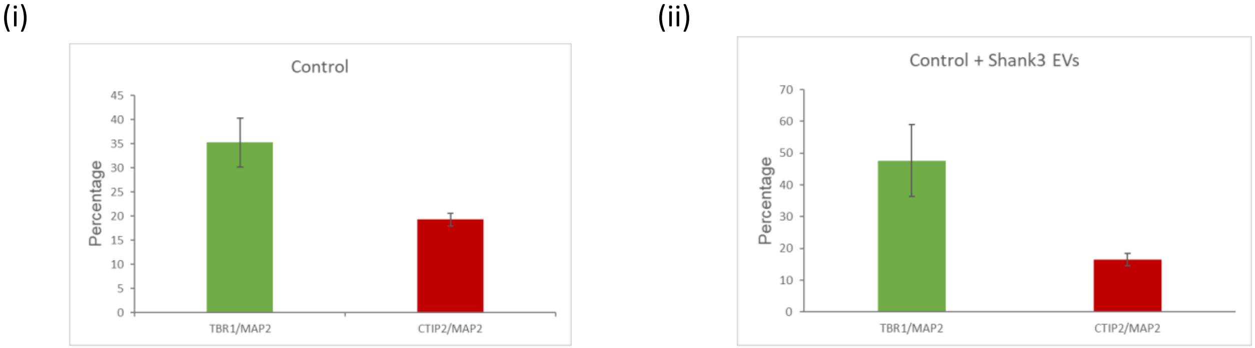

#### Supplementary Figure 2

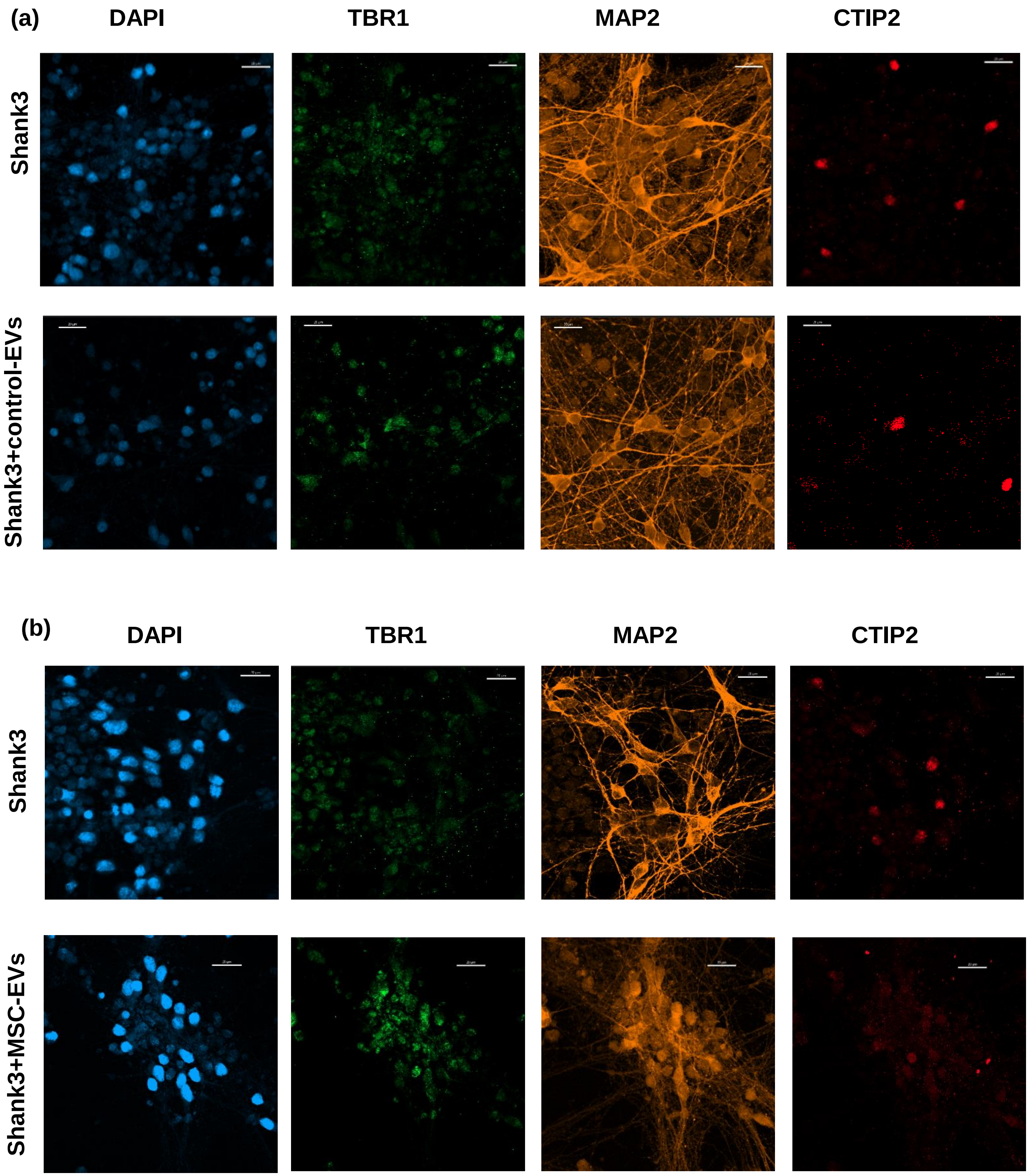

### Supplementary Figure 3

(a)

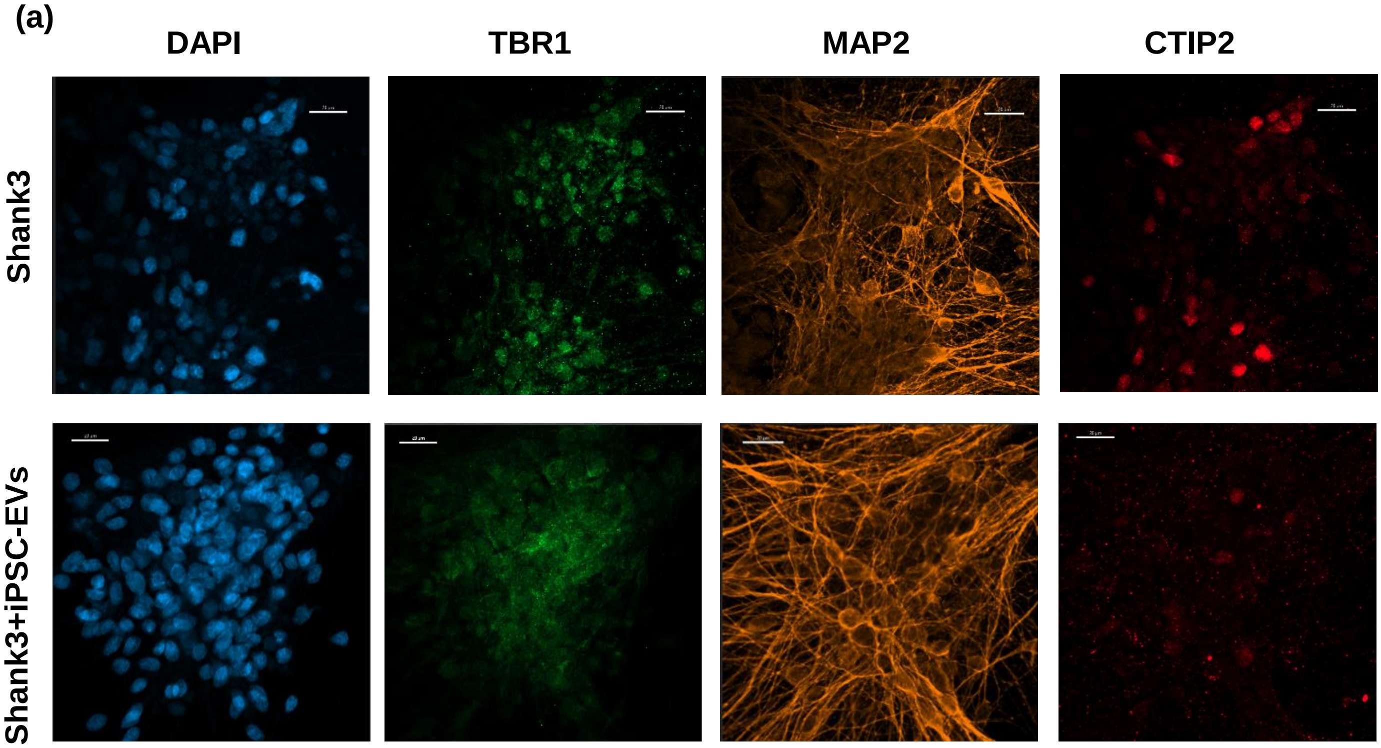


(b)

(i)

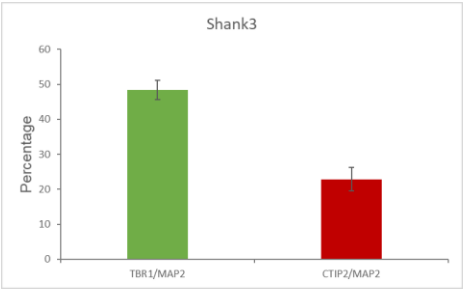


(ii)

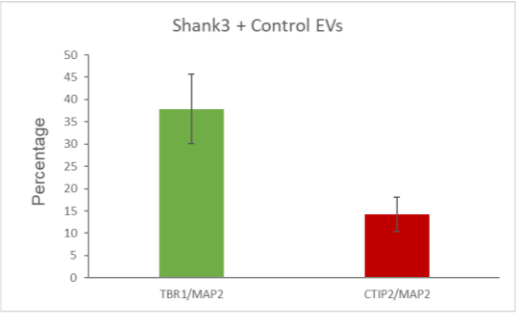


(iii)

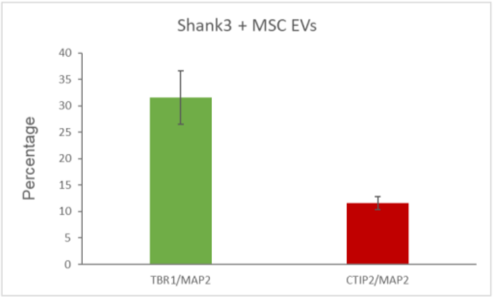


(iv)

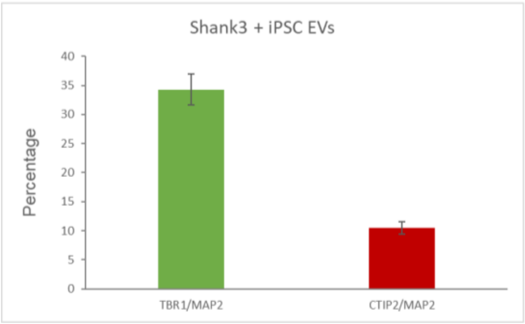

### Supplementary Figure 4

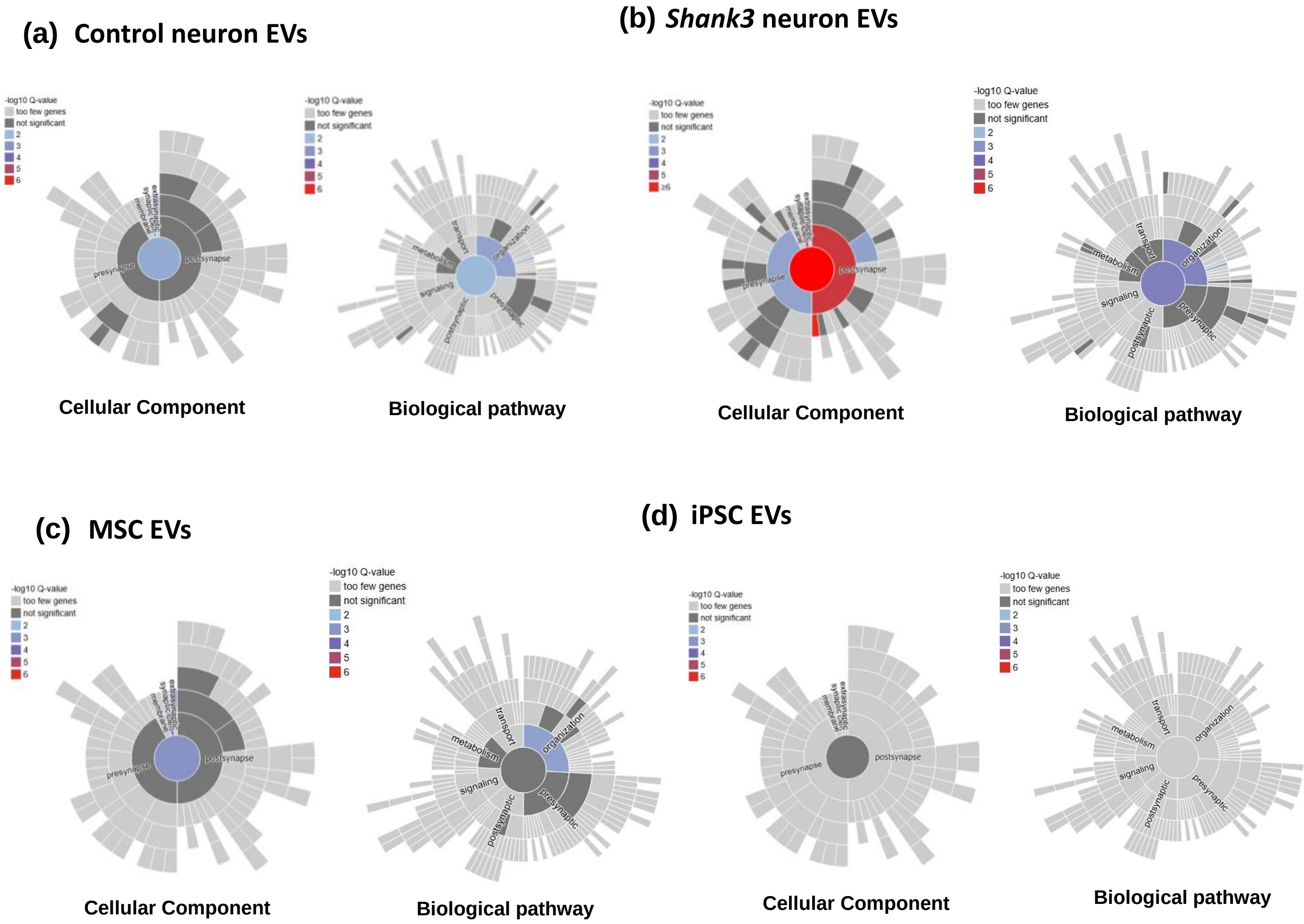

### Supplementary Figure 5

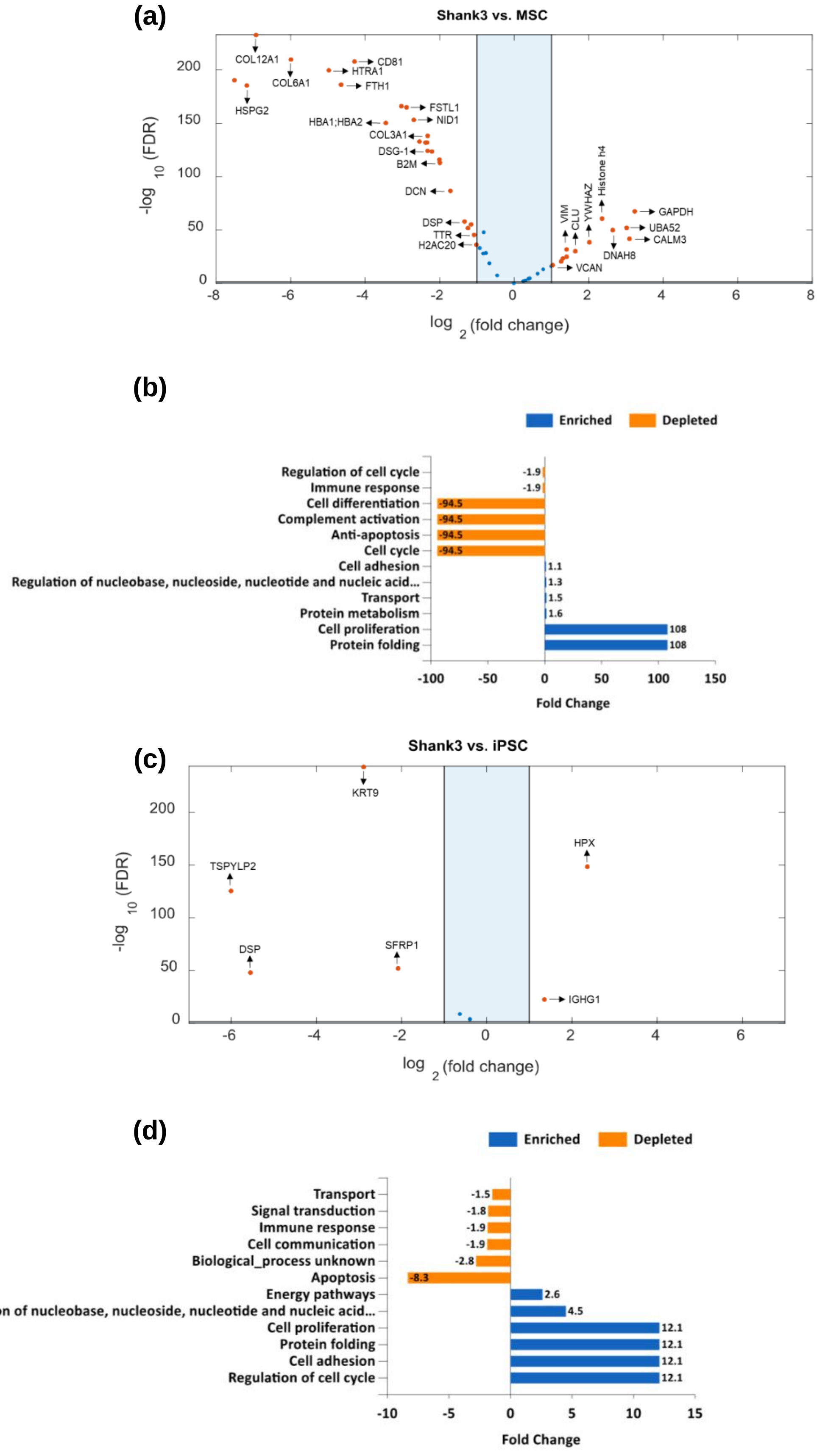
